# KMT2A modulates the epigenetic landscape of rDNA by facilitating the recruitment of histone lysine acetyltransferase PCAF to the rDNA locus

**DOI:** 10.64898/2026.09.02.747590

**Authors:** Kaisar Ahmad Lone, Anita Choudhary, Ajay Kumar Mahato, Shweta Tyagi

## Abstract

Histone acetylation is often associated with transcriptional activation across a wide range of genes, playing a key role in RNA Polymerase II dynamics. However, its specific role in transcriptional activation of RNA Polymerase I (RNA Pol I) remains unclear. In this study, we demonstrate that KMT2A associates with ribosomal DNA (rDNA) loci. Notably, the loss of KMT2A in our inducible KO cell line does not affect the levels of H3K4me3 on rDNA. We support these observations with analyses of ChIP-seq data from mouse embryonic stem cells. While H3K4 methylation remains unchanged, the absence of KMT2A causes a significant decrease in H3 acetylation levels on rDNA, especially in H3K9 acetylation levels. To identify the histone lysine acetyl transferases (KATs) that cooperate with KMT2A in promoting rDNA transcription, we examined the occupancy of multiple KATs and their associated histone acetylation marks on the rDNA locus. Our analyses identify PCAF (p300/CBP associated factor) as the KAT that contributes to KMT2A-mediated transcriptional activation of rDNA. Depletion of KMT2A reduces the levels of PCAF on rDNA, suggesting that KMT2A plays the role of a co-activator of RNA Pol I in the recruitment of KATs on the rDNA loci. Finally, the depletion of KMT2A leads to the disruption of the pre-initiation complex from the rDNA 47S promoter, resulting in a stalled RNA Pol I complex at the spacer promoter. Our findings elucidate the non-redundant, distinct function of KMT2A in the regulation of RNA Pol I transcription.

## Introduction

Ribosomal DNA (rDNA), which encodes ribosomal RNA (rRNA), is transcribed into the 47S rRNA precursor by a distinct transcription machinery comprising RNA Polymerase I (RNA Pol I) and various co-regulatory factors within the nucleolus. RNA Pol I drives the transcription of rDNA, producing nascent 47S rRNA, which is subsequently processed into the different functional rRNA components of the ribosome. This process must be tightly coordinated with the synthesis of 5S rRNA by RNA Pol III and the production of ribosomal protein transcripts by RNA Pol II. Such coordination is achieved through the interplay of various cis-regulatory elements and trans-acting factors that modulate transcription. While the number of identified trans-acting factors is relatively limited in RNA Pol I transcription, the process initiates with the formation of the pre-initiation complex (PIC) at both the extensively studied 47S ribosomal DNA (rDNA) core promoter and the less characterised Spacer promoter (Sp). This process involves the stepwise recruitment of multiple accessory proteins, culminating in the assembly of the RNA Pol I complex. The Upstream Binding Factor (UBF) plays a crucial, multifaceted role not only in PIC assembly but also in promoter escape and transcription elongation by facilitating the recruitment of additional transcriptional components (Sharifi & Bierhoff, 2018; Goodfellow & Zomerdijk, 2013; De Ponti et al., 2025). A central player in PIC formation is the Selectivity Factor 1 (SL1), a multi-subunit complex comprising the TATA-binding protein (TBP) and four associated transcription factors: TAF1A/TAFI48, TAF1B/TAFI63, TAF1C/TAFI110, and TAF1D/TAFI41 (Learned et al., 1985; Eberhard et al., 1993; Gorski et al., 2007; Comai et al., 1994). SL1 interacts with the acidic C-terminal region of the UBF and thus stabilizes the transcriptional machinery at the promoter and enables the recruitment of the 14-subunit RNA Pol I complex through the Pol I-associated factor RRN3 (Comai et al., 2004). The final formation of the PIC is achieved when this ternary complex comprising UBF, SL1, and RNA Pol I assembles at the rDNA promoter, establishing a transcriptionally competent state (Friedrich et al., 2005).

rDNA transcription is regulated at multiple levels, including post-translational modifications of histones within rDNA-associated nucleosomes and DNA methylation at CpG sites. The impact of DNA methylation on transcription depends on its genomic context within the rDNA locus. Specifically, methylation at the promoter region suppresses rDNA transcription, whereas methylation within the gene body region has been shown to enhance transcriptional activity (Huang et al., 2021). Another level of regulation of rDNA transcription is through the regulation of the histone proteins. Various epigenetic modifiers introduce distinct modifications to these histone proteins, which can either facilitate rDNA transcription by making the chromatin accessible or repress it by maintaining a closed chromatin state. The precise mechanisms by which various epigenetic modifiers, in particular the H3K4 histone methyltransferases regulate rDNA transcription remain an open question, warranting further investigation. H3K4 methylation is deposited by the lysine methyltransferase 2 (KMT2)/COMPASS/Mixed Lineage Leukemia (MLL) protein families. In mammals, the COMPASS family comprises six members: MLL1-4/KMT2A-D, SET1A-B/KMT2F-G. These proteins function within complexes containing four common subunits, collectively known as WRAD (WDR5, RBBP5, ASH2L, and DPY30) (Sugeedha et al., 2021). The KMT2 family primarily regulates transcription through the Su (var)3-9, Enhancer-of-zeste, Trithorax (SET) domain. Additionally, certain members, such as KMT2A and KMT2B, possess a transcriptional activation domain (TAD), further contributing to their regulatory roles (Cosgrove & Patel, 2010). KMT2A is the founding member of this family and is involved in the progression of leukemia through its gene translocations. One unique feature of KMT2A is the recruitment of lysine histone acetyltransferases (KATs) through its TA domain (Ernst et al., 2001). Indeed, the p300/CBP, PCAF (p300/CBP associated factor) and MOF, which acetylate the various lysine residues in histone H3 and H4, have been shown to associate with KMT2A to activate transcription (Ernst et al., 2001; Dou et al., 2005; Revenko et al., 2010; Goto et. al, 2002). H3K9 acetylation plays an important role in mediating the switch from transcription initiation to transcriptional elongation by facilitating the recruitment of the super elongation complex on the chromatin (Gates et al., 2017). However, the role played by H3K9 acetylation in the regulation of RNA Pol I transcription is not known.

Recently, we have reported that the depletion of KMT2A leads to a significant reduction in rDNA transcription levels (Lone et al., 2026). Transcript analysis and extensive complementation assays using mutations of KMT2A attributed this function to its TA domain. Additionally, through various biochemical approaches, we established that KMT2A interacts with the RNA Pol I machinery (Lone et al., 2026). Nonetheless, the mechanistic details through which KMT2A utilizes its TA domain to regulate rRNA transcription remained unclear. Here, we show that KMT2A binds to rDNA loci across multiple cell lines. Interestingly, KMT2A depletion does not alter H3K4me3 levels on rDNA, a finding corroborated by analyses of ChIP-seq data from mouse embryonic stem cells (mESC). Despite the unchanged H3K4 methylation levels, KMT2A depletion leads to a dramatic reduction in H3 acetylation levels on rDNA, particularly H3K9 acetylation. We profiled the binding of various KATs on rDNA and used inducible KO cell lines (iKOs) to demonstrate that KMT2A recruits PCAF to rDNA. The depletion of KMT2A results in the disruption of the pre-initiation complex at the rDNA 47S promoter, causing the RNA Pol I complex to stall at the spacer promoter. Our findings suggest that KMT2A-recruited KATs are requisite for the transition of RNA Pol I to productive transcription from the 47S core promoter. Taken together, our results demonstrate how KMT2 enzymes can regulate rDNA loci by distinct pathways.

## Results

### KMT2A binds to the ribosomal DNA (rDNA) locus

Recently, we have shown that KMT2 enzymes bind to rDNA loci in human cells by ChIP-seq analyses (Lone et al., 2026). The quantitative analysis using primer coordinates confirmed that ChIP-qPCR data corresponded with the ChIP-seq data analyses (Lone et al., 2026). *However, as this binding will be the basis of our entire study, we initiated the process by validating our prior analyses through the execution of new ChIP experiments.* We used select primers that span the rDNA unit in its entirety as follows: RNA Pol I promoter, including Spacer and core 47S promoter (Primer #1-3), the transcribed gene (Primer # 4-8) and the intergenic spacer (IGS) region, which is not transcribed by RNA Pol I (Primer #9-18). HoxA9, a canonical RNA Pol II-transcribed gene, was used as a reference for KMT2A binding. CD4 promoter, which does not show KMT2A binding, was used as a negative control (Fig. 1A). Our results reveal that KMT2A shows enriched binding at both the RNA Pol I promoters: the spacer promoter and the 47S promoter. We also observed higher binding in the IGS (Fig. 1A).

**Figure 1.**
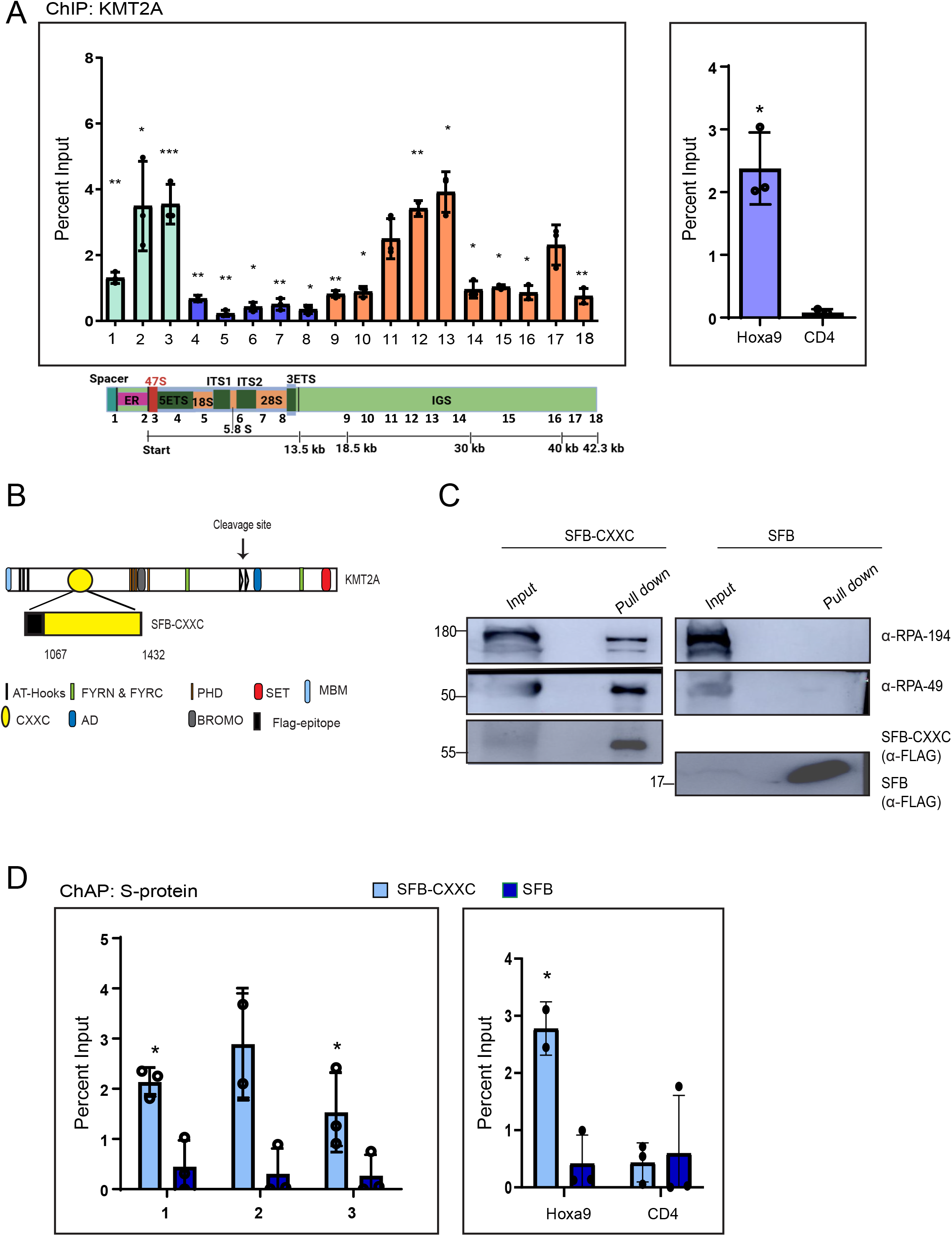
KMT2A binds to the rDNA locus through the CXXC domain. **A.** This figure presents the ChIP analysis of KMT2A in HEK293 cells, with HoxA9 used as a positive control for KMT2A binding and CD4 as a negative control. Various primer sets targeting human rDNA were used to assess KMT2A binding across the rDNA locus (indicated in the schematic below. Also see Supplementary Table 1for primer coordinates). Primer sets 1–3 (green) target the promoter regions, primers 4–8 (blue) correspond to the transcribed region of the RNA Pol I, and primers 9–18 are specific to the Intergenic spacer (IGS) region of rDNA. Data represented as means ± SD from three individual biological replicates. Significance for each rDNA primer was calculated in comparison with the negative control CD4. Error bars represent SD. \**P* ≤ 0.05, ** *P* ≤ 0.005, \*\*\**P*≤ 0.0005, \*\*\*\**P*≤ 0.00005, ns: not significant *P* > 0.05 (two-tailed Students *t* test). **B-C.** A schematic of the full-length KMT2A construct, with its various domains, highlighting the CXXC domain fragment, spanning amino acids 1067 to 1432, is shown. (**B**). Ectopically expressed SFB-CXXC and SFB alone were subjected to S-protein pull-down assays (**C**). The CXXC pull-down was detected using an anti-FLAG antibody, and the immunoblot was probed with antibodies against RPA194 and RPA49 to assess the interaction of the CXXC domain with different components of the RNA Pol I complex. SFB, used as a mock control, was also detected with the anti-FLAG antibody. Molecular weight marker is indicated on the left. **D**. Chromatin affinity purification (ChAP) analysis of CXXC and the SFB vector control is shown. Data are presented as per cent input. The significance of binding for each primer was determined by comparing the per cent input of CXXC binding relative to the vector control (SFB). Error bars indicate standard deviation (SD). Results are shown as mean ± SD from three independent biological replicates. Statistical significance was assessed using a two-tailed Student’s t-test, with *P ≤ 0.05 indicating significance and ns (not significant) for P > 0.05.

In order to understand if this binding was cell-line specific or extended to other cells, we used previously published KMT2A ChIP-seq data from mESCs and mapped it to the custom mm39*-*rDNA v1.0 assembly. In this assembly, a single unit of 45kb rDNA (mouse) called Chromosome R has been added (George et al., 2023). Similarly, we obtained data for H3K4me2 and H3K4me3 in the mESCs and aligned them using the same genome assembly. As shown in Figure S1A, KMT2A bound to the entire region of the rDNA sequence, including the promoter and transcribed unit of RNA Pol I and the IGS. In contrast to KMT2A binding, H3K4me2 and H3K4me3 showed discrete peaks in the IGS of the rDNA with high enrichment at the promoter and transcribed unit of RNA Pol I (Fig. S1A-B).

We next performed ChIP in mouse embryonic fibroblasts (MEFs) to assess KMT2A binding across different regions of the mouse rDNA repeat using a specific primer set (Fig. S1B-C, Zentner et al., 2014). To ensure that we score specific binding, we included an IgG ChIP as a mock control. qRT-PCR analysis revealed that KMT2A binds at the spacer promoter, enhancer repeat, and 47S promoter regions of mouse rDNA. Additionally, we observed KMT2A binding within the transcribed region and the IGS (Fig. S1B-C). The consistent binding of KMT2A across different cell lines here suggests that its association with rDNA is conserved across species.

KMT2A recruitment to the chromatin is governed either by the direct binding of KMT2A to the chromatin through its CXXC domain or through the ternary complex of MLL-MENIN-LEDGF (Birke et al; 2002, Yokoyama et al; 2008, Caslini et al; 2007,). MLL contains the MENIN binding motif (MBM), MENIN interacts with the LEDGF, a chromatin-associated protein, and this ternary complex in turn docks KMT2A on the chromatin (Yokoyama et al; 2008). However, the CXXC domain of KMT2A plays an important role in the direct recruitment to the non-methylated CpG islands of the chromatin (Ayton et al; 2004, Erfurth et al; 2008). At the same time, the CXXC domain plays an important role in the tethering of the Polymerase associated factor complex (PAFc), an RNA Pol II-associated factor, to KMT2A. PAF complex is implicated in RNA Pol II-driven transcription, where it regulates the rate of transcription elongation (Krogan et al., 2003, Sims et al., 2004). The interaction of KMT2A with the PAFc was mapped to the CXXC domain of KMT2A extending from 1067-1432 amino acids (Milne et al., 2010). Based on our observations so far, we predicted that the RNA Pol I complex might be interacting with KMT2A in a similar fashion. To test our hypothesis, we cloned the CXXC fragment of KMT2A (1067-1432 amino acids) into the SFB (S protein, FLAG epitope, and streptavidin-binding peptide) containing vector and performed the pull-down studies. We transiently expressed SFB-CXXC and subjected the cells to S-protein pull-down. SFB (alone) expressing cells were used for mock pull-down. RNA Pol I is a 14-subunit holo-complex, out of which 7 subunits are unique (Viktorovskaya et al., 2015). Previously, we have shown that KMT2A interacts with various subunits of the RNA Pol I complex, SL-1 complex, UBF and RRN3 (Lone et al., 2026). Here, we probed for two unique RNA Pol I subunits: RPA-194, the largest subunit, and RPA-49, a peripheral subunit involved in initiation and elongation (Kuhn et al., 2007) (Fig. 1B-C). Our results indicate that the CXXC was able to pull down both subunits successfully, indicating a positive interaction.

To further demonstrate that the CXXC domain targets KMT2A to the rDNA promoter, we performed chromatin-based affinity purification (ChAP) using S-protein beads (Zargar et al., 2018). This was conducted in control cells expressing the SFB-tagged vector and experimental cells expressing SFB-CXXC. Our results revealed a significant enrichment of CXXC domain on the rDNA promoter compared to the control cells (Fig 1D). Taken together, our results suggest that the CXXC domain may play a crucial role in facilitating KMT2A interaction with the RNA Pol I complex and its binding to the rDNA locus.

### KMT2A depletion does not alter H3K4me3 levels on the rDNA locus

We have previously shown that H3K4me2 and H3K4me3 marks decorate the rDNA, and they are required for RNA Pol I transcription (Lone et al., 2026). Here, we have observed that KMT2A binds the rDNA promoter (Fig. 1). To confirm how KMT2A influences the status of H3K4 methylation, we used inducible knockouts of KMT2A (iKO) in human cell lines, HEK293 (Fig. 2A) (Malik et al., 2023; Chinchole et al., 2022). We observed that KMT2A levels were substantially reduced on the promoter, as well as the transcribed region of rDNA loci in KMT2A iKO cells (Fig 2A-D; *Note to the Reviewers: The results presented in Fig. 2 are exclusive to this manuscript and are not part of Lone et al., 2026*). Despite this marked decrease in the KMT2A levels, H3K4me3 levels were reduced only at the Spacer promoter and remained unchanged elsewhere on the rDNA locus. Similarly, H3K4me2 levels were reduced only at the promoter region (Primer#1-3; Fig. 2A-D). In contrast, the H3K4me3 and H3K4me2 levels were significantly decreased along with KMT2A levels on the HoxA9 gene, indicating that our knockout induction had worked (Fig. S2A-C). Taken together, our findings reveal that KMT2A regulates rDNA transcription without altering H3K4me3 marks at the rDNA locus in human cells.

**Figure 2.**
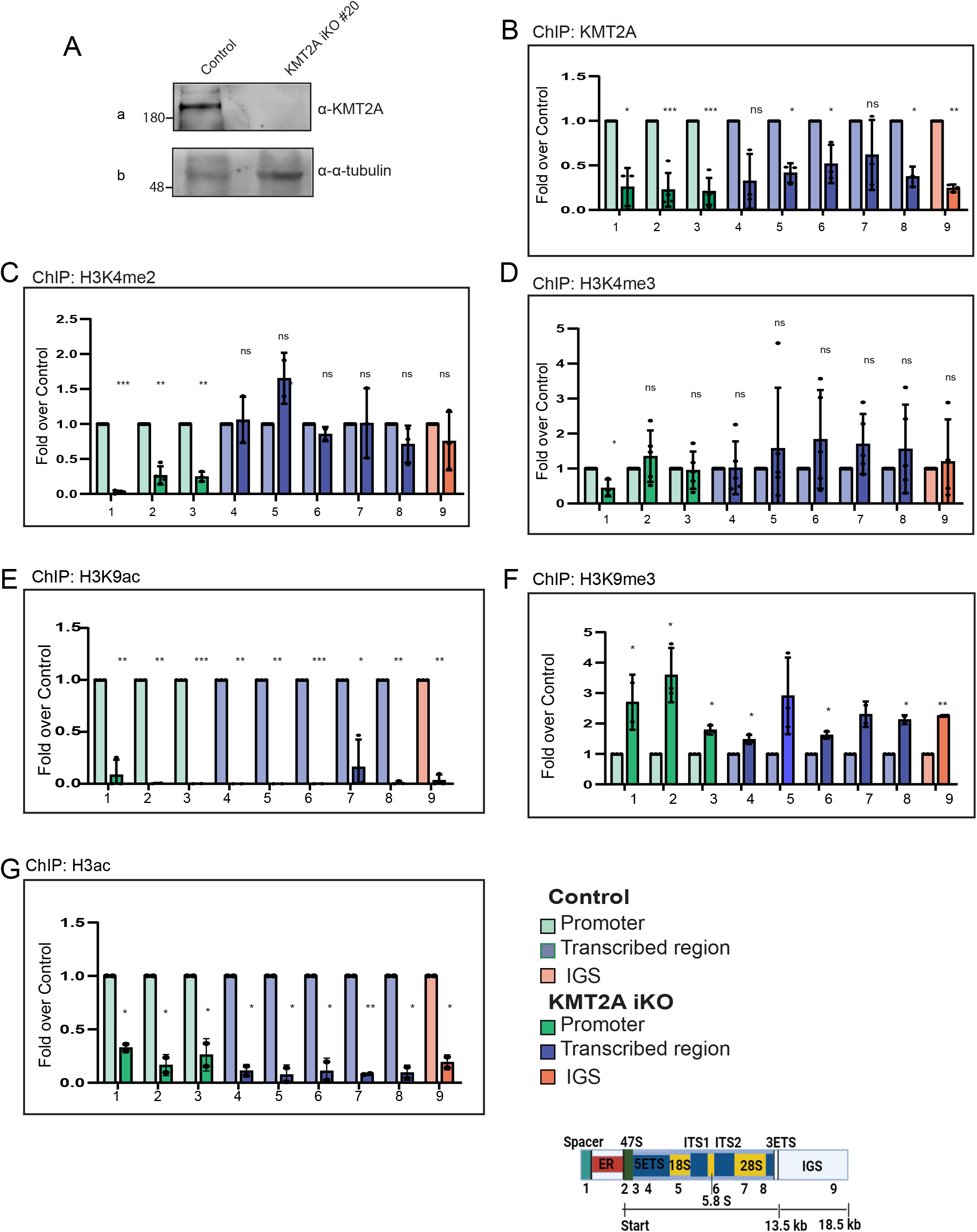
Effect of the KMT2A depletion on the epigenome of the rDNA locus. **A**. Immunoblot shows KMT2A protein levels in CRISPR-Cas9-generated inducible knockout cells (iKO#20). Blots were probed with α-KMT2A and α-α-tubulin as shown. **B-D.** ChIP analyses of KMT2A (**B**), H3K4me2 (**C**), H3K4me3 (**D**), H3K9ac (**E**), H3K9me3 (**F**), and H3ac (**G**) in HEK293 cells in control and KMT2A iKOs are shown. Primers 1–9 were used to assess KMT2A and histone modification binding across the human rDNA locus as described above. Error bars represent the mean ± SD from three independent experiments for all except the H3ac (two biological replicates). Statistical significance between control and test conditions was determined using a two-tailed Student’s t-test (*P ≤ 0.05, **P ≤ 0.005, ***P ≤ 0.0005).

In order to confirm our results further, we analysed previously published data where inhibition of KMT2A activity was achieved by treating *Mll1^fl/fl^* mESCs*; Cre-ER^TM^* ESCs with either 4-hydroxytamoxifen (4-OHT) for *kmt2A deletion* (KMT2A mKO) or use of MM-401, which specifically blocks KMT2A enzymatic activity by disrupting its interaction with WDR5 (KMT2Ai) (Zhang et al., 2019; Cao et al., 2014). A comparison between the control, KMT2Ai, or KMT2A mKO revealed that there was no detectable change in the H3K4me3 levels on the RNA Pol I transcribed rDNA (Fig. S2D). However, we observed a slight decrease in the discrete peaks of IGS and on the spacer promoter (Fig. S2D). When analysed, we observed a decrease in levels of H3K4me2 on the promoter region but no change in the IGS region of the rDNA (Fig. S2E). To validate our findings here, we analyzed canonical target genes from previous data sets, which reported reduced H3K4me3 levels in KMT2A KO MEFs (Wang et al., 2009). Our analysis revealed a reduction in H3K4me3 levels at select HoxA clusters, including HoxA3 and HoxA10 genes (Fig. S2F), thereby supporting the validity of our analysis pipeline.

### KMT2A modulates the epigenetic landscape of the rDNA via histone acetylation

Even though we did not observe any change in H3K4me3 levels on rDNA upon loss of KMT2A, rRNA transcription is affected (Lone et al., 2026). To understand what may impede transcription from this locus, we examined H3K9 acetylation (H3K9ac), another mark linked to open chromatin in KMT2A iKO cells. To our surprise, we found that H3K9ac levels were drastically reduced throughout the rDNA locus as well as on the canonical target genes like HOXA9 but not U2_C_, an upstream region from *U2* snRNA gene used as a negative control here (Fig. 2E, S2G, Malik et al., 2023). This decrease in H3K9ac levels was accompanied by a 2-fold increase in the H3K9me3 levels on rDNA as well as on HOXA9, while U2_C_ showed no significant change (Fig. 2F, S2H). As active transcription can also influence the state of histone acetylation (Martin et al., 2021), we performed ChIP with an antibody recognising pan H3 acetylation in KMT2A iKO (Fig. 2G, S2I). Curiously, the H3ac was reduced on the promoter region and in the transcribed region of the rDNA, as well as on the canonical target gene HOXA9. Our results here indicate that the role of the KMT2A complex is to enhance an open chromatin state by promoting H3 acetylation for successful rRNA transcription.

### Screening for the different H3 and H4 acetylation marks on rDNA locus

Multiple KATs, including PCAF, GCN5, CBP, p300, MOF, MOZ and TIP60, have been implicated in transcriptional activation through the deposition of distinct histone acetylation marks. However, the specific histone acetylation signatures associated with active rDNA transcription remain poorly defined. Since our data indicated that KMT2A promotes rDNA transcription via histone acetylation, we sought to determine which histone acetylation marks and their associated KATs are preferentially enriched across the human rDNA locus and therefore might cooperate with KMT2A in regulating rDNA transcription.

To systematically address this question, we analysed publicly available ChIP-seq datasets of canonical histone H3 and H4 acetylation marks on lysine residues generated in HEK293 cells. Among these, H3K9ac was prioritized because this mark acts in direct opposition to the repressive H3K9 methylation mark, and establishes an open chromatin environment conducive to transcription initiation and elongation (Bian et al., 2011 ). To our knowledge, only one H3K9ac dataset in HEK293 cells was available, although used as a control in the study with FLAG overexpression (Weber et al., 2023). The raw sequencing reads were reanalysed by mapping them to the rDNA region in customized human hg38-rDNA reference genome. Our analysis revealed robust H3K9ac enrichment across the entire rDNA repeat, with the strongest enrichment observed at the rDNA promoter and in the IGS regions (Fig. 3).

**Figure 3.**
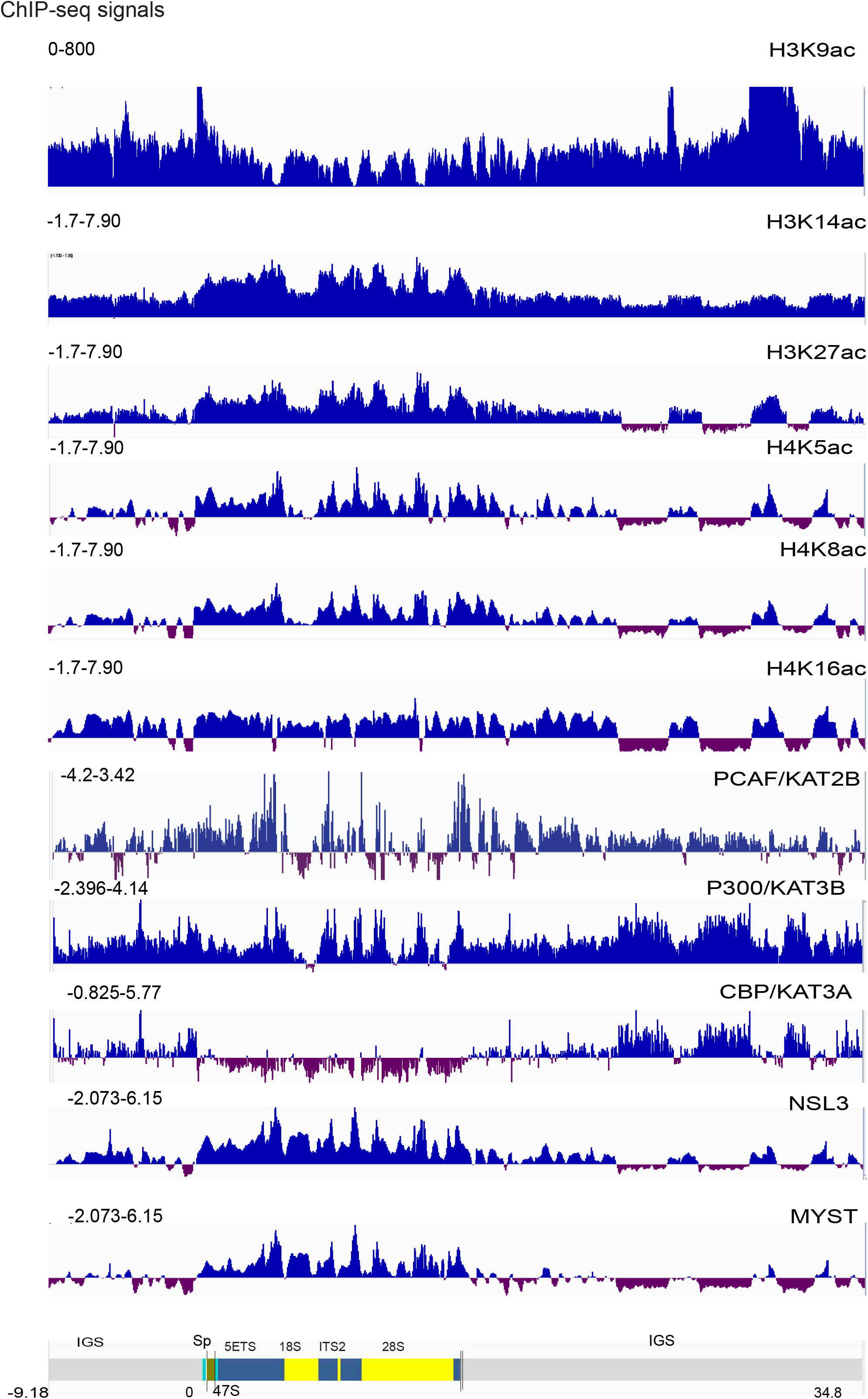
Binding profiles of histone H3 and H4 acetylation and their associated lysine acetyltransferases (KATs) on human rDNA locus. The ChIP-seq analysis of the various histone acetylation: H3K9ac, H3K14ac, H3K27ac, H4K5ac, H4K8ac & H4K16ac and the associated KATs (PCAF, P300, CBP, NSL3 and MOF) are presented as enrichment over input. *Note: the H3K9ac track shown here is generated from cells overexpressing the FLAG tag construct and Y axis is truncated at 800 for better visualization. The complete profile is provided in Supplementary Figure S3K.* All ChIP-seq tracks are from HEK293 cells, except PCAF, which was generated from HCT116 cells.

Since multiple histone acetylation marks are associated with active chromatin, particularly H3K14ac and H3K27ac, we next investigated the genome-wide distribution of these modifications at the rDNA locus (Savitsky et al., 2016). Both H3K14ac and H3K27ac were significantly enriched across the rDNA repeat (Fig. 3), further supporting the notion that actively transcribed rDNA exists within a highly acetylated chromatin environment.

The H4K16 histone acetyltransferase MOF has been shown to functionally cooperate with the KMT2A complex, with both chromatin modifiers co-occupying transcriptionally active promoters to establish an open chromatin environment that facilitates gene expression (Dou et al., 2005; Wang et al., 2023). Moreover, MOF-dependent H4K16 acetylation is indispensable for the maintenance of KMT2A-driven transcriptional programs, highlighting the close functional relationship between KMT2A-mediated H3K4 methylation and H4 acetylation (Valerio et al., 2017). Based on these observations, we hypothesised that KMT2A-dependent regulation of rDNA transcription may also be associated with the deposition of multiple H4 acetylation marks. To investigate this possibility, we analysed publicly available ChIP-seq datasets for H4K5ac, H4K8ac, and H4K16ac after remapping the sequencing reads to our customized hg38-rDNA reference genome (Liu et al., 2022). Interestingly, all three acetylation marks displayed highly similar distribution profiles across the rDNA repeat, with pronounced enrichment throughout the transcribed region and comparatively discrete enrichment within the IGS (Fig. 3). Together, these findings indicate that the human rDNA locus harbours a complex histone acetylation landscape comprising both H3 and H4 acetylation marks. Given the established functional interplay between KMT2A-mediated transcriptional activation and several histone acetyltransferase complexes, our data support a model in which KMT2A coordinates the establishment of an active chromatin environment at rDNA through the recruitment or stabilization of one or multiple KAT activities.

### Profiling the binding of lysine acetyltransferases on the rDNA

Following the establishment of the histone acetylation landscape across the rDNA locus, we next sought to identify the KATs responsible for depositing these acetylation marks. To this end, we analysed publicly available ChIP-seq datasets of different KATs to determine their occupancy across the rDNA. KATs are classified into three major families based on the structural organization of their catalytic domains and their evolutionary origins: (i) GNAT family, comprising GCN5 (KAT2A) and PCAF (KAT2B); (ii) p300/CBP family, comprising p300 (KAT3B) and CBP (KAT3A); and (iii) MYST family, comprising TIP60 (KAT5), MOZ (KAT6A), HBO1 (KAT7), and MOF (KAT8). To investigate the GNAT family, we analysed the ChIP-seq dataset of PCAF in HCT116 cells (Perea-Resa et al., 2017), as no suitable PCAF or GCN5 ChIP-seq dataset was available for HEK293 cells. Our analysis revealed that PCAF is enriched across the rDNA locus, with prominent discrete peaks within the coding region and a relatively continuous distribution throughout the IGS (Fig. 3). Next, we examined the binding profiles of the p300/CBP family(Byun and Gardner, 2013), where both p300 and CBP displayed discrete binding patterns across the rDNA. Interestingly, CBP exhibited little or no enrichment within the coding region, whereas p300 showed occupancy in both the coding and intergenic regions (Fig. 3), suggesting differential recruitment and distribution of these two closely related acetyltransferases on rDNA. To investigate the MYST family, we first analysed ChIP-seq data for KANSL3 (NSL3) (Liu et al., 2022). KANSL3 is a core component of the NSL (Non-Specific Lethal) complex, which consists of MOF (KAT8), KANSL1 (NSL1), KANSL2 (NSL2), KANSL3 (NSL3), MCRS1, PHF20, WDR5, and HCF1. KANSL3 functions as a scaffold protein that facilitates recruitment of MOF to chromatin. Therefore, KANSL3 occupancy was used as an indirect measure of MOF recruitment to the rDNA. Analysis of the KANSL3 ChIP-seq data revealed strong enrichment across the rDNA coding region, whereas only discrete peaks were observed within the IGS (Fig. 3). To validate these observations, we further analysed an independent ChIP-seq dataset generated using a MOF (MYST1/KAT8) antibody. The MOF ChIP-seq profile closely resembled that obtained with KANSL3, displaying strong enrichment throughout the coding region of the rDNA. Although a few peaks within the IGS were absent in the MOF dataset, the overall binding pattern remained highly consistent between the two datasets (Fig. 3), supporting the conclusion that MOF is recruited to both the coding and intergenic regions of the rDNA. Finally, we investigated the binding of MOZ (KAT6A), another member of the MYST family, using ChIP-seq data generated with FLAG-tagged MOZ (Weber et al., 2023). MOZ displayed pronounced enrichment throughout the coding region of the rDNA together with a distinct peak within the IGS (Fig. S3K). Collectively, these analyses demonstrate that members of all three major KAT families occupy the rDNA locus, although each exhibits a distinct binding profile. These observations suggest that multiple acetyltransferases contribute to the establishment and maintenance of the rDNA histone acetylation landscape. Having established the occupancy of these KATs on rDNA, the next objective was to determine which of these acetyltransferases are recruited in a KMT2A-dependent manner and therefore may function downstream of KMT2A to regulate rDNA transcription.

### Identification of KATs Involved in the Regulation of rDNA Transcription

To determine which of the identified KATs are functionally involved in the regulation of rDNA transcription, we individually depleted the KATs that were found to occupy the rDNA locus (discussed above) using shRNA-mediated knockdown. Specifically, MOF, MOZ, p300, CBP, PCAF, and GCN5 were targeted. Transcript analysis confirmed efficient depletion of each KAT (Fig.S3A). Upon transcription, the 13 kb 47S precursor ribosomal RNA (pre-rRNA) undergoes processing at its 5’ end, known as the 5’ external transcribed spacer (5’ETS) (McStay & Grummt, 2008). Consequently, it serves as an indicator of pre-rRNA transcript levels using quantitative real-time PCR (qRT-PCR). We assessed rDNA transcription by quantifying the levels of the 5′ETS. Individual depletion of each KAT resulted in a significant reduction in 5′ETS transcript levels (Fig. S3B), indicating that all of these acetyltransferases contribute to the maintenance of rDNA transcription. To further evaluate the relative contribution of these KATs to rDNA transcription, we performed gain-of-function studies by transiently overexpressing KATs for which cDNA was available— p300, CBP, PCAF, and GCN5— in HEK293 cells. (*Mammalian expression constructs for MOF and MOZ were not available to us and, therefore, could not be included in this analysis.*) qRT-PCR analysis of the 5′ETS transcript revealed that overexpression of all four KATs enhanced rDNA transcription to varying extents. Notably, PCAF overexpression resulted in the most pronounced increase in 5′ETS transcript levels compared with p300, CBP, and GCN5 (Fig.S3C-J), suggesting that PCAF exerts a stronger stimulatory effect on rDNA transcription than the other KATs examined. Taken together, these findings demonstrate that multiple KATs positively regulate rDNA transcription, with PCAF exhibiting the strongest transcriptional activation upon overexpression, suggesting that it may be a key regulator of RNA Pol I-mediated rDNA transcription.

### MYST family KATs might be regulating the rDNA transcription indirectly

As described above, we observed the occupancy of the MYST family KATs: MOF and MOZ, at the rDNA locus. Since both proteins have established roles in transcriptional regulation and are functionally associated with KMT2A, we next investigated whether they directly regulate rDNA transcription. We first focused on MOZ by analysing the published ChIP-seq datasets generated from HEK293 cells overexpressing FLAG-tagged wild-type MOZ, and a catalytically inactive KAT mutant (Weber et al., 2023). As expected, the KAT mutant did not exhibit a significant reduction in MOZ occupancy across the rDNA locus compared with the wild-type protein. These findings suggest that MOZ association with rDNA is independent of its catalytic activity. To further examine the contribution of MOZ-mediated histone acetylation, we analysed H3K9ac ChIP-seq datasets from cells expressing the vector control (FLAG), wild-type MOZ or the KAT mutant (Weber et al., 2023). Genome-wide comparison of the rDNA locus revealed only minor differences in H3K9ac enrichment among these conditions, with modest changes confined to a few regions around the rDNA promoter and IGS (Fig.S3K). No substantial loss of H3K9 acetylation was observed upon disruption of MOZ catalytic activity. Together, these observations further support the notion that MOZ influences rDNA transcription indirectly rather than through direct acetylation of rDNA-associated chromatin.

As MOF is also enriched at the rDNA locus, we next investigated whether MOF directly regulates rDNA transcription. To address this, we analysed published MOF and histone H4 acetylation ChIP-seq datasets generated from mESCs in which MOF was acutely depleted using an auxin-inducible degron system (Erdogdu et al., 2026). Analysis of MOF occupancy confirmed its association with the mouse rDNA locus under control conditions, with a remarkable reduction following auxin treatment (Fig.S3L). Although the overall binding pattern differed somewhat from that observed in HEK293 cells (in the enzyme and H4 acetylation described marks below), these differences are likely attributable to species-specific and developmental differences between mESCs and HEK293 cells. Because MOF catalyses acetylation of multiple lysine residues on histone H4, we examined the distribution of H4K16ac, H4K12ac, H4K8ac and H4K5ac following acute MOF depletion. Surprisingly, none of these histone H4 acetylation marks exhibited a significant reduction across the rDNA locus after MOF depletion (Fig.S3L). Collectively, these analyses indicate that neither MOZ nor MOF directly regulates rDNA transcription through local histone acetylation at the rDNA locus. Instead, both MYST family members are more likely to influence rDNA transcription indirectly, possibly through regulation of other chromatin-associated factors or transcriptional networks. Based on our observations here, we decided not to pursue further mechanistic studies on the MYST family or the p300/CBP family of acetyltransferases. Instead, we focused on the GNAT family, where PCAF emerged as our strongest choice as (i) its overexpression produced the most pronounced elevation in rDNA transcription among all KATs examined here, (ii) PCAF has been shown to acetylate H3K9 and H4 residues on the rDNA locus (Shen et al., 2013), and (iii) it has been implicated in activating transcription in association with KMT2A (Revenko et al., 2010). These observations prompted us to investigate whether PCAF mediates rDNA transcription via KMT2A.

### KMT2A recruits the lysine acetyltransferase PCAF to the rDNA locus

Our previous RNAi experiments, followed by complementation analyses, showed that the KMT2A TA domain, but not its SET domain, was required for the transcription of 47S rRNA (Lone et al., 2026). To further validate these findings, we overexpressed KMT2A and KMT2A mutants—KMT2AΔSET (a point mutation N3906A, which renders the SET domain catalytically inactive, Patel et al., 2009) and KMT2AΔTAD in HEK293 cells. When we assessed 5′ETS transcript levels under Control (vector alone), KMT2A and mutant overexpression conditions, our results show that 5′ETS transcript levels showed a 2-fold increase upon expression of KMT2A (Fig. S4A-B). This increase was observed with the expression of KMT2AΔSET also. However, cells expressing KMT2AΔTAD displayed no increase in 5’ETS transcript and behaved like the Control sample, despite comparable KMT2A expression (Fig. S4A-B). Previously, we reported SET- and TA domain-mediated regulation of alpha-satellite transcripts by KMT2A, which are also non-coding transcripts (Malik et al., 2023). To validate our findings, we used alpha-satellite transcripts as a control. Unlike wild-type KMT2A, neither KMT2AΔSET nor KMT2AΔTAD mutant expression could increase alpha-satellite transcript levels (Fig. S4C), thus further confirming the specificity of the TA domain-mediated role of KMT2A in the regulation of the rDNA transcription.

To investigate this further, we interrogated the association of KMT2A with PCAF, and our endogenous IP with KMT2A revealed that it can co-immunoprecipitate endogenous PCAF robustly and specifically (Fig. S4D). We then used a truncation of MLL, which has the TA domain (KMT2A D1, Chodisetty et al., 2024), and could successfully pull down PCAF using it, but not from the vector with the same epitope tags (Fig. S4E), indicating that the TA domain of KMT2A was sufficient to interact with PCAF. PCAF was shown to bind the rDNA promoter previously(Shen et al., 2013). We confirmed the association of PCAF with the promoter and transcribed region of rDNA by ChIP (Fig. 4A). Next, we performed PCAF ChIP in KMT2A iKOs. Consistent with our hypothesis that KMT2A recruits KATs, we observed a significant reduction in PCAF binding to the rDNA promoter region, as well as in the transcribed unit (Fig. 4B, S4F). Taken together, our findings indicate that KMT2A recruits lysine acetyltransferase, PCAF, to the rDNA locus to promote open chromatin configuration and transcriptional activation.

**Figure 4.**
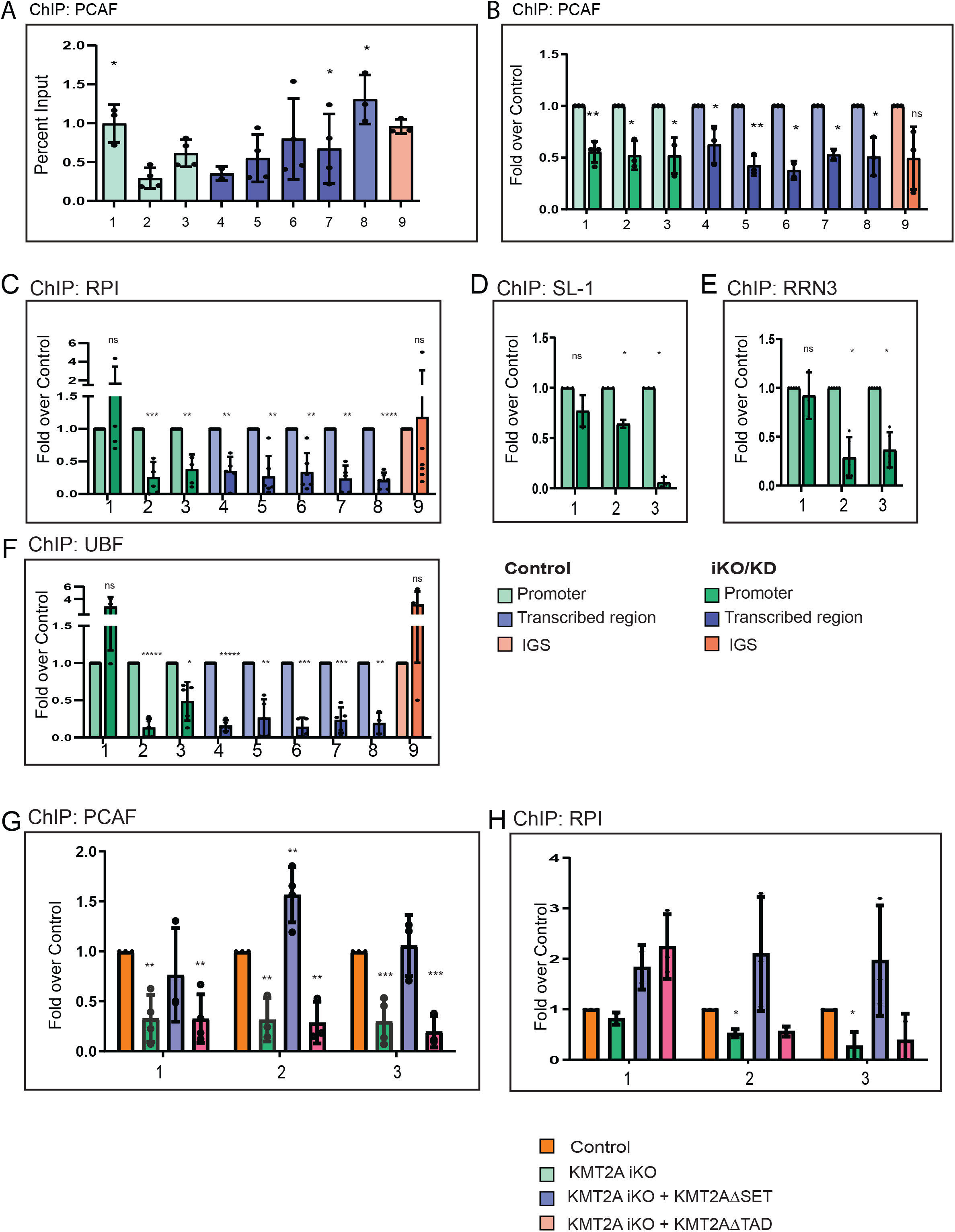
KMT2A recruits PCAF to promote RNA Pol I Pre-Initiation Complex Assembly at the rDNA Locus. **A.** ChIP analysis of PCAF binding on rDNA in HEK293 cells is shown. Data are presented as per cent input from three individual biological replicates. Error bars represent the mean ± SD. **B-F.** ChIP analyses of PCAF (**B**) and various RNA Pol I components involved in pre-initiation complex formation at the rDNA locus, including RNA Pol I (**C**), SL1 (**D**), RRN3 (**E**), and UBF (**F**) are shown. Primers 1–3 were used to assess the binding of the promoter-associated factors, SL1 and RRN3, while primers 1–9 were used to examine the binding of PCAF, RNA Pol I and UBF in both control and iKO cells. Data are presented from three or more biological replicates. Error bars represent the mean ± SD. Statistical significance was determined using a two-tailed Student’s t-test (*P ≤ 0.05, **P ≤ 0.005, ***P ≤ 0.0005, ****P ≤ 0.00005; ns: not significant, P > 0.05). **G-H.** shows the PCAF and RNA Pol I ChIP in the KMT2A iKO cells exogenously expressing the KMT2AΔSET or KMT2AΔTAD. 1, 2 & 3 primers were used to assess binding on the spacer promoter, enhancer repeats and 47S promoter, respectively. Error bars represent the mean ± SD. Statistical significance was determined using a two-way ANOVA between the control and each test condition. (*P ≤ 0.05, **P ≤ 0.005, ***P ≤ 0.0005; ns: not significant, P > 0.05).

### Loss of KMT2A abrogates RNA Pol I pre-initiation complex formation on 47S promoter

Loss of KMT2A resulted in decreased rRNA transcription without affecting the H3K4me3 status, but dramatically reducing the H3 acetylation. To determine exactly how this rearranged epigenome impacts the RNA Pol I transcription, we performed additional experiments. In the human rDNA transcription process, UBF initially activates the rDNA by binding to the RNA Pol I promoter, followed by recruiting the SL1 complex to form the PIC at the rDNA promoter (Panov et al., 2006). UBF then recruits RNA Pol I through interactions with its RPA34 and RPA49 subunits, whereas SL1 associates with RRN3. These factors collectively engage the 47S and spacer promoters, thereby facilitating the initiation of transcription at the rDNA locus. Once the formation of the PIC stabilises RNA Pol I, RRN3 is released from the RNA Pol I complex, allowing RNA Pol I to continue with transcript elongation and produce 47S pre-rRNA (Grummt, 2003; McStay & Grummt, 2008).

Upon loss of KMT2A, we observed reduced binding of RNA Pol I at the 47S promoter (and transcribed unit) but not at the spacer promoter (Fig. 4C). The same observations were made for SL-1 complex and RRN3 (Fig. 4D-E). UBF binding also remains intact at the spacer promoter, but it is reduced on the 47S promoter (Fig. 4F). We have shown using comparative transcriptomic analysis previously that loss of KMT2A does not globally disrupt the transcriptional programs governing ribosome biogenesis, rRNA processing, or ribosomal protein production (Lone et al., 2026; Malik et al., 2023). Our findings here indicate that the PIC assembles at the spacer promoter, but it is unable to progress to the core promoter in the absence of KMT2A. It has been suggested that the spacer promoter functions as a high affinity binding site or an entry site for RNA Pol I as well as UBF (De Winter & Moss, 1986). Our results suggest that KMT2A facilitates the transfer of RNA Pol I PIC from the spacer promoter to the 47S promoter once it has assembled at the spacer promoter.

### Loss of KMT2A results in a stalled RNA Pol I complex at the spacer promoter

Our ChIP experiments in KMT2A iKO cells indicate that loss of KMT2A may result in a stalled RNA Pol I complex at the spacer promoter. The spacer promoter has been shown to produce transcripts, which terminate at a promoter-proximal terminator (Kuhn & Grummt, 1987). Additionally, UBF has been proposed to promote RNA Pol I promoter escape by facilitating the transition of RNA Pol I complex that produces short abortive transcripts from the 47S promoter to a stable elongation complex capable of producing full-length transcripts (Goodfellow & Zomerdijk, 2013). To verify that the RNA Pol I complex, which was assembled at the spacer promoter, is transcriptionally competent, we tested for transcripts originating from the spacer promoter. As shown, upon knockdown of KMT2A, transcripts could be detected from the spacer promoter (primer #1) but not from or after 5’ETS region (primer #3, 3’, 3’’), indicating that the stalled RNA Pol I complex could form PIC at spacer promoter but could not progress to productive transcription of rRNA from 47S promoter in absence of KMT2A (Fig S4G-H). Our results indicate that KMT2A acts as a co-activator of RNA Pol I to ensure that it can progress to the core promoter to perform productive transcription.

### The KMT2A-TA domain-mediated recruitment of PCAF ensures progression of RNA Pol I to the 47S promoter for productive transcription

Taken together, we propose that KMT2A facilitates the transfer of RNA Pol I PIC from the spacer promoter to the 47S promoter following its assembly at the spacer promoter by recruiting PCAF to rDNA through its TA domain. To further strengthen this hypothesis, we performed rescue experiments in KMT2A iKO cells. Briefly, KMT2A knockout cells were transfected with either the KMT2A ΔSET or KMT2A ΔTAD, as described above. We then assessed the occupancy of PCAF and RNA Pol I on the rDNA promoter by ChIP analysis under control and rescue conditions. Our results showed that exogenous expression of KMT2A ΔSET efficiently restored the recruitment of both PCAF and RNA Pol I to the rDNA promoter. In contrast, expression of the KMT2AΔTAD mutant failed to rescue the occupancy of either PCAF or RNA Pol I (Fig. 4G–H, S4I). These findings provide strong functional evidence that the TA domain of KMT2A is essential for PCAF recruitment to rDNA chromatin. Furthermore, the concomitant restoration of RNA Pol I occupancy by KMT2A ΔSET, but not by KMT2A ΔTAD, indicates that TA domain-dependent recruitment of PCAF is a critical step in establishing a transcriptionally competent rDNA promoter and promoting efficient rDNA transcription.

## Discussion

### KMT2A acts as a co-activator of RNA Pol I by recruiting PCAF to the rDNA locus

Here, we have shown that KMT2A binds to rDNA loci in human and mouse cells. Our data suggest that KMT2A is responsible for the transition of RNA Pol I from the spacer promoter to the core promoter. Interestingly, loss of KMT2A does not affect the tri-methylation levels of H3K4 but is accompanied by a drastic reduction in H3K9ac as well as pan H3 acetylation. This decrease in acetylation comes with an increase in H3K9 methylation.

A distinctive characteristic of KMT2A is its ability to recruit KATs via its TA domain, especially CBP/P300. Collaborative interactions between CBP, the KMT2A TA domain and various transcriptional activators have been shown to achieve synergistic boosts in transcriptional activation (Goto et al., 2002).

Generally, KATs like MOF, MOZ and HBO1 have also been reported to be associated with KMT2A to activate transcription (Dou et al., 2005, Takahashi et al., 2021). In our previous study, we showed that KMT2A utilises its TA domain to activate rRNA transcription. Here we reinforced our hypothesis by 5’ETS transcript analyses in cells overexpressing KMT2A or its TA domain deletion. We further show that KMT2A interacts with PCAF, a KAT not previously established as a common KMT2A-associated factor and recruits it to rDNA. Taken together, our results implicate KMT2A in the recruitment of various KATs to the rDNA to keep it open for rDNA transcription.

Our observations here align with those made in yeast, where it was shown that H3K4 and H3K36 methylation play an important role in the recruitment of the NuA3 a histone lysine acetyltransferase complex on the promoters of transcriptionally active genes (Martin et al., 2017). Other chromatin remodelling complex like B-WICH are also reported to play an important role in the recruitment of various KATs on the rDNA locus, as the depletion of B-WICH results in reduced levels of H3-acetylation and particularly the H3K9-acetylation levels in human cell lines (Vintermist et al., 2011). Interestingly, PCAF has also been shown to acetylate a subunit of the SL-1 complex, thereby enhancing its DNA-binding activity and promoting rDNA transcription (Muth et al., 2001). Taken together with our findings, these observations raise the possibility that KMT2A-mediated histone acetylation and TTF-I– mediated acetylation of the SL-1 complex represent distinct steps in the regulation of rDNA transcription. Alternatively, TTF-I may function as a component of a larger regulatory KMT2A/PCAF/SL-1 complex. Future studies will be required to determine how these factors are functionally coordinated at the rDNA locus.

Histone acetylation plays an important role in the transcriptional activation of the rDNA gene clusters. CBP and two other histone acetyltransferases, namely p300 and PCAF, were found to stimulate rDNA transcription when overexpressed in NIH 3T3 cells (Hirschler-Laszkiewicz et al., 2001). Similarly, TIP60, a histone acetyltransferase, has been identified on the rDNA promoter, where it activates UBF by acetylating it, thereby promoting the transcriptional activation of rDNA (Halkidou et al., 2004). Although the repertoire of KATs has been identified on rDNA, the precise mechanisms governing their recruitment to the rDNA locus and their mechanistic details in the regulation of the rDNA transcription require further validation. Our results provide an insight into how large chromatin modifying complexes like KMT2A may ensure the recruitment of KATs to the rDNA locus to activate transcription.

### Impact of rDNA transcription in KMT2 rearranged leukemia

Elevated ribosomal transcription by RNA Pol I is one of the important hallmarks of cancer (Barna et al., 2008). In a recent development, the significance of rRNA biogenesis in fostering malignant characteristics was highlighted by the observation that leukemic cells exhibit a heightened dependence on increased rDNA transcription, rendering them highly sensitive to RNA Pol I inhibition (Bywater et al., 2012). KMT2 members, particularly KMT2A rearrangements, play a crucial role in causing cancer/leukemia. KMT2A commonly undergoes rearrangement with various partner genes, leading to the formation of KMT2A-fusion proteins. These chimeric proteins lead to elevated transcript levels observed in both Acute myeloid leukemia (AML) and Acute lymphoid leukemia (ALL). It has been proposed that the principal activity contributing to the oncogenic nature of KMT2A-fusion proteins is the TBP (part of SL1 complex) loading activity (Okuda et al., 2016). It is important to note that a recent study demonstrated use of a selective inhibitor of RNA Pol I transcription, CX-5461, showed effective treatment outcomes for aggressive AML, including KMT2A-driven AML, surpassing standard chemotherapies (Hein et al., 2017). Aberrant recruitment of KATs and other effector proteins has also been proposed to drive the oncogenicity of MLL-fusion proteins (Ernst et al., 2025). However, whether such complexes can deregulate RNA Pol I-dependent transcription remains unexplored. Taken together, these studies indicate the necessity for deeper investigation into how KMT2A-fusion proteins deregulate the transcription of the rDNA locus in disease conditions, while our study firmly establishes their regulatory role in RNA Pol I transcription.

## Funding

K.A.L. is a recipient of Junior and Senior Research Fellowships of Council of Scientific and Industrial Research, India, toward the pursuit of a PhD degree of Regional Centre for Biotechnology. A.C is a recipient of Junior Research Fellowship of the University Grant Commission, India, toward the pursuit of a PhD degree of Regional Centre for Biotechnology. This work was supported in part by DBT Wellcome Trust India Alliance Senior Fellowship to S.T.[IA/S/18/2/503981] and CDFD core funds.

## Acknowledgments

We are thankful to Geethanjali Ravindran for cloning the CXXC domain of KMT2A in the pcDNA-SFB vector and Lakshana Laali for helping in the shRNA cloning of KATs.

## Author Contributions

K.A.L. conducted all experiments and ChIP-seq analysis. A.K.M further helped in developing the pipeline for rDNA ChIP-seq analysis. A.C. performed the experiments shown in Supplementary Fig. 3A-J. S.T. and K.A.L. designed the experiments. K.A.L and. S.T. wrote the manuscript.

## Conflict of Interest

The authors declare that they have no conflict of interest.

## Data availability statement

All data generated or analyzed during this study are included in the manuscript and supporting files; source data files have been provided for all figures.

## Materials and Methods

### Cell culture

HEK293 (human embryonic kidney) and Mouse embryonic fibroblasts (MEFs) cells were cultured in DMEM (Thermo Fisher Scientific) supplemented with 10% (v/v) fetal bovine serum (FBS), 1% (v/v) GlutaMAX, and 100 U/ml penicillin-streptomycin. Cell line authentication was done via short tandem repeat (STR) profiling (Life Technologies). Cells were maintained at 37°C in a humidified atmosphere with 5% CO .

### Cloning, stable cell line generation, and transfections

To generate the CXXC domain clone of KMT2A, the CXXC fragment was amplified from the pBabe-FLAG KMT2A full-length construct using Phusion polymerase and subcloned into the XhoI site of the pcDNA-SFB vector. The C subunit of KMT2A cloned into the pBabe-FLAG vector was used to generate various KMT2A mutants. The KMT2AΔTAD mutant was generated by deleting amino acids 2847–2855 using site-directed mutagenesis. Additionally, a point mutation (N3906A) was introduced to generate the KMT2AΔSET mutant. The KMT2A_C_D1 (referred to as MLL_C_ D1 previously) fragment was generated by cloning the KMT2A fragment (2718-3280 amino acids) in the pcDNA-SFB vector in XhoI site.

KMT2A inducible knockouts were generated in HEK293 cells as previously described (Chinchole et al., 2022; Malik et al., 2023). For RNA-based experiments, endogenous KMT2A protein levels were depleted using RNA interference (RNAi), following established protocols (Malik et al., 2023). Briefly, siRNA duplexes were designed and transfected using Oligofectamine (Thermo Fisher Scientific), with a siRNA sequence targeting the firefly luciferase gene serving as a control. Cells underwent two rounds of transfection at 24-hour intervals and were harvested 72 hours post-transfection. For the transfection of the CXXC in HEK293 cells, cells were transfected with the SFB-CXXC and the SFB vector control separately with PEI and harvested after 36 hrs, then subjected to the Co-immunoprecipitation experiments or for the ChAP (Chromatin-affinity purification). For the overexpression of KMT2A and its mutants KMT2AΔSET and KMT2AΔTAD in HEK293 cells, the cells were transfected with PEI with these different constructs and harvested after 36 hours and were subjected to RNA isolation. shRNAs targeting PCAF, GCN5, MOF, MOZ, CBP, P300 were designed according to the TRC shRNA design guidelines and cloned into the pLKO.1 vector (Addgene). Briefly, the pLKO.1 vector was digested with AgeI and EcoRI, and the linearized vector was ligated with annealed oligonucleotides encoding different KAT-specific shRNA sequences.

### Co-immunoprecipitation and immunoblot analysis

HEK293 cells transiently transfected with pcDNA-SFB-CXXC, pcDNA-SFB-KMT2A D1 or pcDNA-SFB were lysed using NETN buffer (100 mM NaCl, 20 mM Tris-HCl pH 8.0, 0.5 mM EDTA, and 0.5% Nonidet P-40) for 30 minutes. The lysates were then subjected to sonication for 10 cycles (each cycle: 30 seconds on, 30 seconds off) at medium power using a Diagenode Bioruptor. Following sonication, lysates were centrifuged at high speed at 4°C. Equal amounts of protein from all lysates were used for S-protein pull-down assays. S-protein agarose beads (Sigma, 69704) were added to each lysate and incubated overnight at 4°C on a nutator. The next day, beads were pelleted by centrifugation at 4°C and washed three times with IP wash buffer (50 mM Tris-HCl pH 7.4, 100 mM KCl, and 0.1% Nonidet P-40). The beads were then resuspended in PBS and Laemmli buffer, boiled for 10 minutes, and subjected to SDS-PAGE. Proteins were transferred onto nitrocellulose (Amersham, 10600003) membrane. For the immunoprecipitation of KMT2A same protocol was followed as described before (Lone et.al 2025). Immunoblotting was performed using antibodies against RPA-49 (Sigma, HPA022-527), PCAF (C14G9), KMT2A (Chinchole et al., 2022), RPA194 (Santa Cruz, SC-48385), and FLAG (Sigma, F7425). Blots were visualised using an ImageQuant LAS 500.

### RNA isolation and qRT-PCR

Total RNA was isolated and treated with TURBO DNase (Thermo Fisher Scientific) for 30 minutes at 37°C to remove genomic DNA. RNA purity was ensured with a no-enzyme amplification step before cDNA synthesis using the SuperScript III Reverse Transcriptase kit (Thermo Fisher Scientific). RT-qPCR was performed on the QuantStudio 5 Real-Time PCR System (Applied Biosystems) with the DyNAmo ColorFlash SYBR Green qPCR kit (Thermo Fisher Scientific). Transcript levels were quantified using the 2-ΔΔCt method (Livak & Schmittgen, 2001). Primer sequences are provided in Supplementary Table S1.

### Chromatin immunoprecipitation

Chromatin immunoprecipitation (ChIP) was carried out following established protocols (Malik et al., 2023; Zargar et.al, 2018; Lone et al., 2026) using the following antibodies: KMT2A-C (A300-374A, Bethyl Labs or as described in Chinchole et al., 2022), H3K4me2 (ab32356, Abcam), H3K4me3 (ab8580, Abcam or 07-473 Millipore), H3K9me3 (ab8898, Abcam), H3K9Ac (ab4441, Abcam), RPA194 (SC-48385, Santa Cruz), UBF (SC-13125, Santa Cruz), RRN3 (Ab-112052, Abcam), H3Ac (ab47915, Abcam), PCAF (C14G9), IgG (12-370, Upstate) and TAF1C (A303-698A, Bethyl Labs). The relative occupancy or per cent input of the immunoprecipitated protein at the ribosomal DNA locus was quantified by RT-qPCR, using the formula: 100 × 2^(Ct Input – Ct IP)^, where Ct Input and Ct IP represent the mean threshold cycles for input and specific immunoprecipitation samples, respectively. Fold change over control values was determined by comparing ChIP results in control cells. Primer sequences can be found in Supplementary Table S1.

### Chromatin affinity purification (ChAP)

Chromatin affinity purification was performed similarly to ChIP, with minor modifications. Briefly, SFB vector control-transfected and CXXC-transfected cells were sonicated using a Diagenode Bioruptor to generate DNA fragments averaging 200–500 bp in size. Ten per cent of the total input was collected from both control and test samples, and precleared lysates were incubated overnight at 4°C with S-protein beads. The next day, 50 microliters of S-protein beads were added to each sample and incubated at 4°C on a nutator for three hours. The beads were then sequentially washed with low-salt, high-salt, LiCl, and TE wash buffers, following previously established protocols (Zargar et al., 2018). Finally, the beads underwent simultaneous decross-linking and elution at 65°C for two hours, followed by PCI purification of the eluted products. The relative occupancy or percent input of the immunoprecipitated protein at the ribosomal DNA locus was quantified by qRT-PCR using the formula: 100 × 2(Ct Input – Ct IP), where Ct Input and Ct IP represent the mean threshold cycles for input and specific immunoprecipitation samples, respectively.

### ChIP-seq analysis

ChIP-seq analysis was performed following the previously described method (Lone et al., 2026). Data from previously conducted KMT2A, H3K4me2, and H3K4me3 ChIP-seq in mESCs (GSE107408) (Zhang et al., 2019) and also MOF, H4K5ac, H4K8ac, H4K12Aac, H4K16 ac (GSE271982) (Erdogdu et.al 2026) were utilized to analyze the mouse rDNA locus. Briefly, the customized genome assembly for the mouse rDNA, mm39-rDNA v1.0 (George et al., 2023) was indexed using Bowtie2 (version 2.3.2). Trimmed reads of KMT2A, H3K4me2, and H3K4me3 from control, MM401-treated, and KMT2A knockout cells as well as MOF, H4K5ac, H4K8ac, H4K12ac, H4K16ac in control and MOF knockout conditions, were aligned to the Bowtie2-indexed mm39-rDNA assembly to generate SAM files. SAM files were then converted to BAM format using Samtools, followed by sorting and indexing. BigWig files were generated using deepTools. Finally, experimental (IP) BigWig (BW) files were normalized to input BW files using the BWCompare command line and the results were visualized in IGV (Integrated Genomic Viewer). For human genomic data pertaining to H3K14ac, H3K27ac (GSE81696) (Savitsky et.al 2016), H4K5ac, H4K8ac, H4K16ac, NSL3, MYST1 (MOF) (GSE198645) (Liu et.al; 2022), P300 and CBP (GSE51117) (Byun and Gardner et al., 2013): MOZ, H3K9ac and its mutants (Weber et al., 2023) (GSE206272). The same pipeline for rDNA analysis was performed as in Lone et.al., 2026.

## Notes

### Competing Interest Statement

The authors have declared no competing interest.

