## Supplemental Information for "KMT2A modulates the epigenetic landscape of rDNA by facilitating the recruitment of histone lysine acetyltransferase PCAF to the rDNA locus"

### Supplementary Material

**Supplementary Figure S1: KMT2A binds to the rDNA locus.**

**A.** The ChIP-seq data analysis for KMT2A, H3K4me2, and H3K4me3 in mouse embryonic stem cells (mESCs) on the mouse rDNA is shown. The ChIP-seq signals for each dataset represent the ratio of normalized values to their respective inputs. At the bottom, the annotation of an individual mouse rDNA repeat highlights its various regions, including the intergenic spacer (IGS), spacer promoter (Sp_prom), 47S core promoter, 5' external transcribed sequence (5_ETS), and 3' external transcribed sequence (3_ETS) .

**B**. The schematic of the individual mouse rDNA repeat and the different primers used for the qRT-PCR in ChIP of mouse-derived cells. Primers #1, 2 & 3 represent the Spacer promoter, Enhancer repeat and 47S promoter, respectively. Primers 4-6 fall in the region transcribed by RNA Pol I, while primers 7-9 represent the IGS region of the rDNA. The coordinates and sequence of each primer are presented in Supplementary Table 1. ITS, Internal Transcribed Spacer.

**C**. The ChIP analysis of KMT2A in mouse embryonic fibroblasts (MEFs) on the rDNA locus with different primers as described in **B**. The MEFs were subjected to ChIP with the KMT2A and IgG antibodies. Data is presented as the percentage input. The significance of the binding is calculated with respect to mock control IgG. Each experiment was performed three times. Error bars represent SD. $*P \leq 0.05$, $**P \leq 0.005$, $***P \leq 0.0005$, $****P \leq 0.0001$, ns: not significant, $P > 0.05$ (two-way ANOVA with Šídák multiple comparison test)

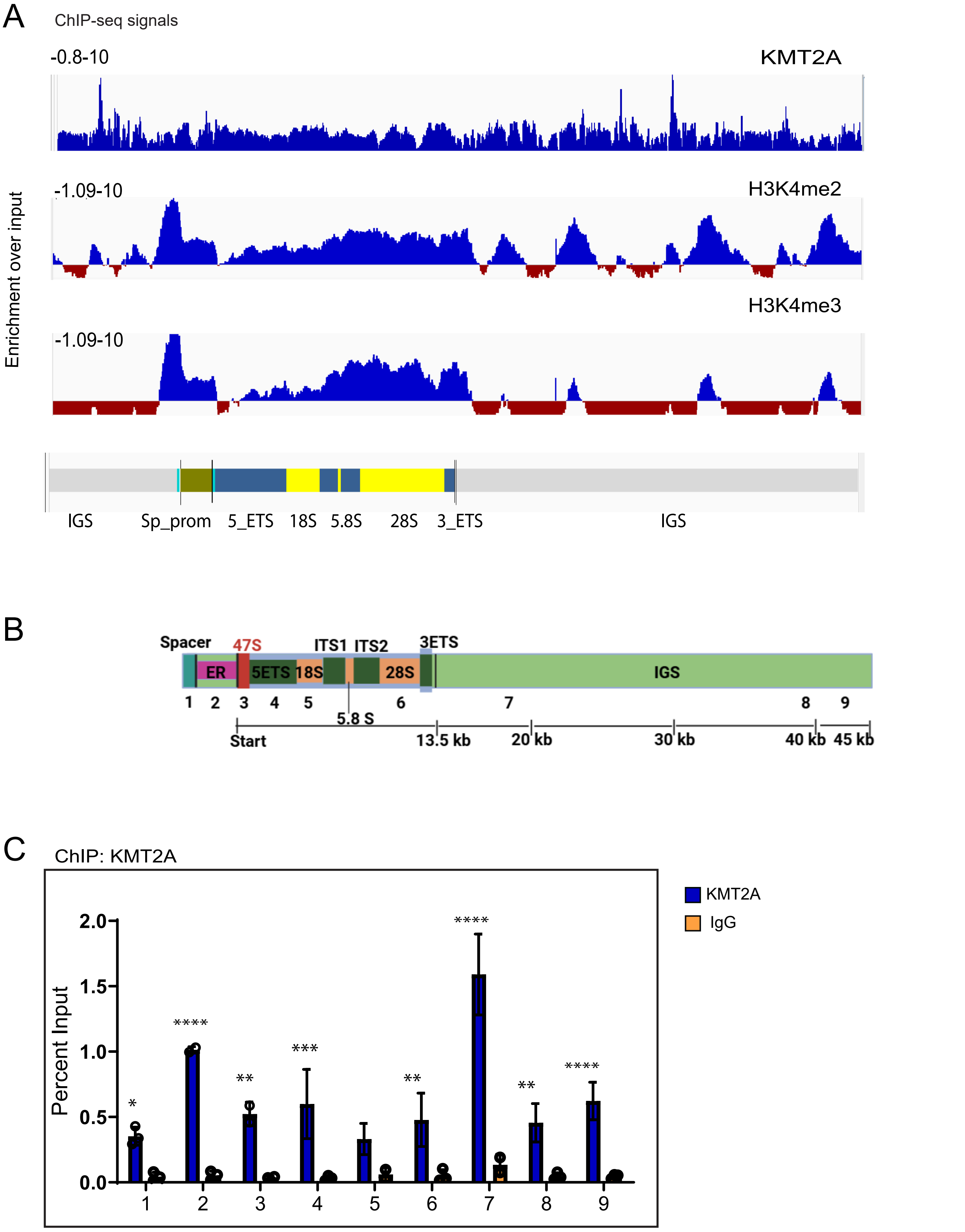


Lone Supplementary Figure 1

**Supplementary Figure S2. Impact of KMT2A depletion on the epigenome of the rDNA locus.**

**A-C.** This figure presents ChIP-seq analysis of KMT2A (**A**), H3K4me3 (**B**) and H3K4me2 (**C**) in control and KMT2A iKO cells. *HoxA9* and *CD4* were used as positive and negative control primers for our ChIP assays shown in Figure 2. Each graph represents data from three independent biological replicates, with error bars indicating standard deviation. Statistical significance was determined using Student's t-test. (*P ≤ 0.05, **P ≤ 0.005, ***P ≤ 0.0005, ****P ≤ 0.00005; ns: not significant, P > 0.05).

**D-E.** ChIP-seq tracks for H3K4me3 (**D**) and H3K4me2 (**E**) enrichment on the mouse rDNA locus under different treatment conditions in mESCs is shown (Zhang et al., 2019, Cao et al., 2014). The top track represents H3K4me3 enrichment in control conditions, while the middle track shows H3K4me3 ChIP-seq data from mESCs treated with the MM401 inhibitor (KMT2Ai), which disrupts the interaction between KMT2A and WDR5. The bottom track represents H3K4me3 ChIP-seq data under KMT2A knockout conditions (KMT2AmKO). The annotation of the mouse rDNA repeat is provided below.

**F.** The ChIP-seq signals of H3K4me3 on the canonical *Hox A* cluster genes in the control, KMT2A inhibition (KMT2Ai), and the KMT2A mKO conditions are shown. *HoxA3* and *HoxA10* regions were further zoomed in to highlight the change in the H3K4me3 levels in the different conditions.

**G-I.** Represents the ChIP analyses of H3K9ac (**G**) H3K9me3 (**H**) H3ac (**I**) in the control and the KMT2A iKO conditions. *HoxA9* and *U2C* were used as the positive and negative control primers for the above-mentioned histone modifications. Data are derived from three or more biological replicates, with error bars representing the mean ± SD. Statistical significance was assessed using a two-tailed Student's t-test (*$P \leq 0.05$, **$P \leq 0.005$, ns: not significant, $P > 0.05$).

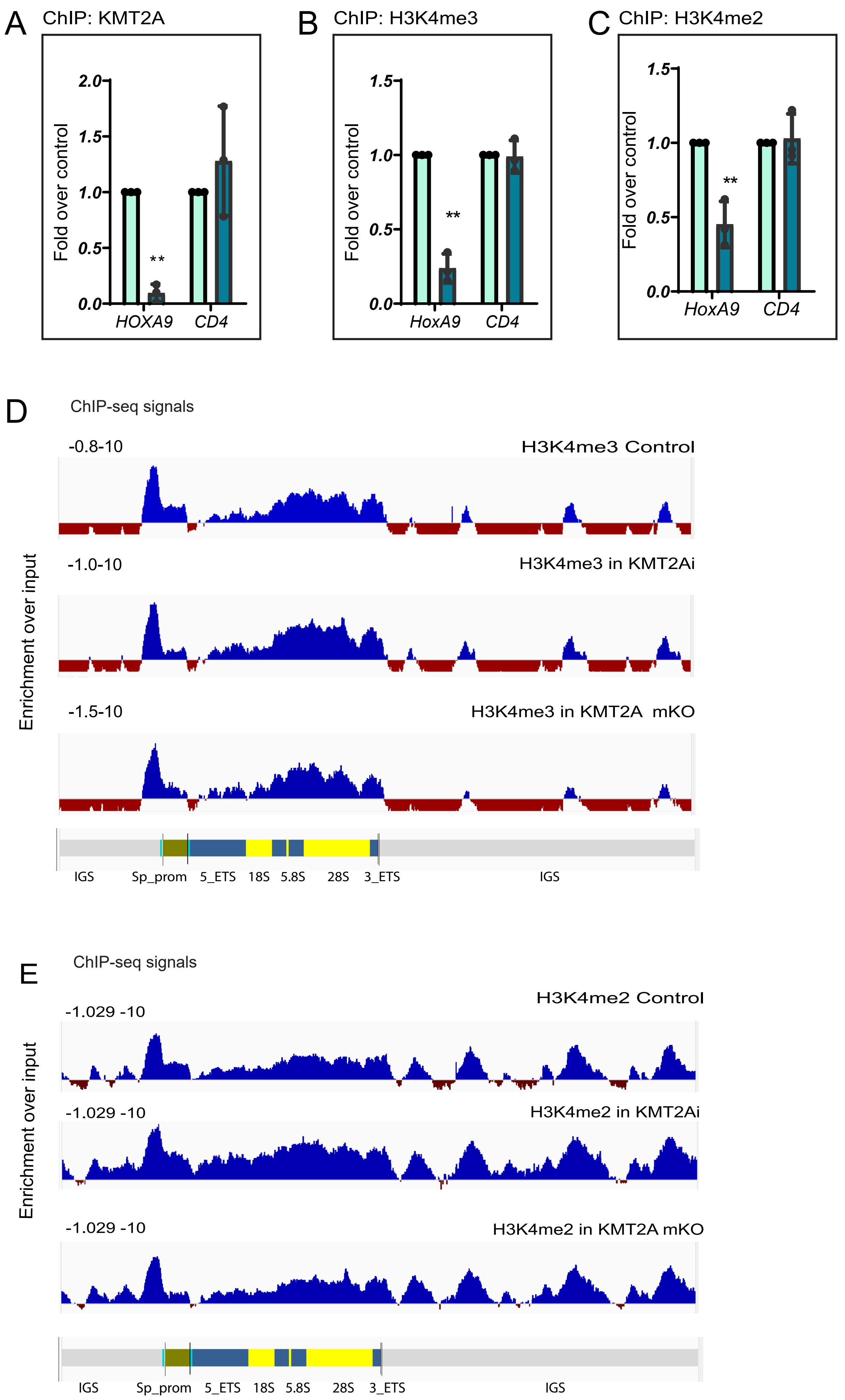


Lone Supplementary Figure 2a

F ChIP-seq signals

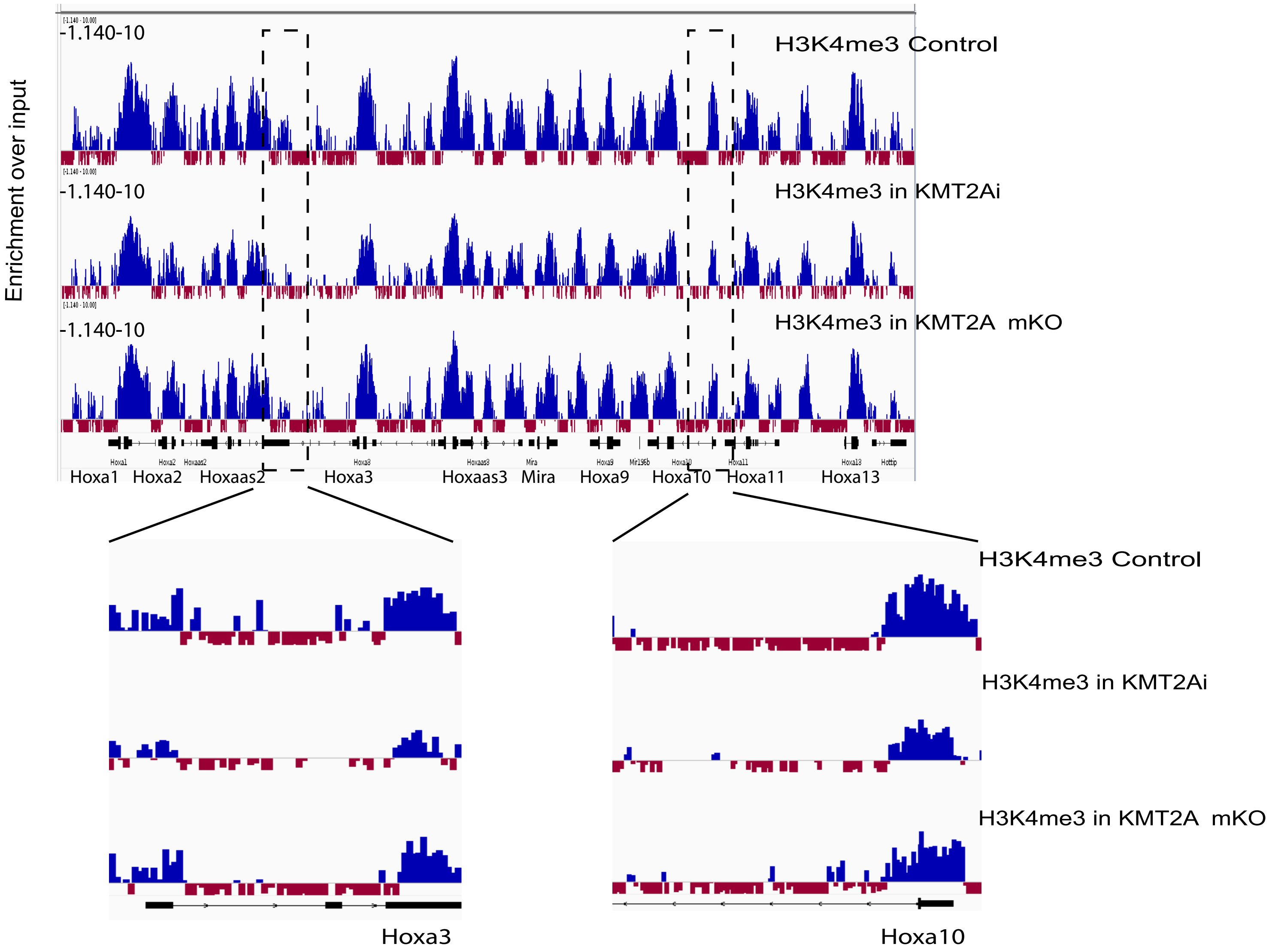


G ChIP: H3K9ac

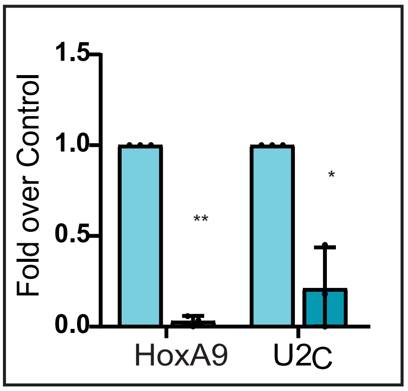


H ChIP: H3K9me3

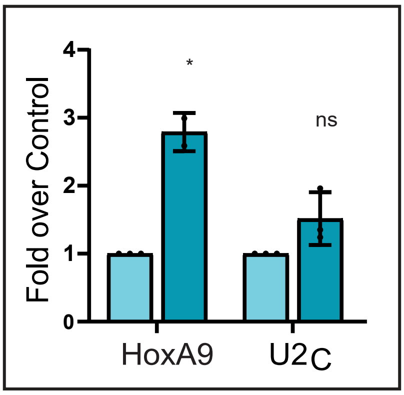


I ChIP: H3ac

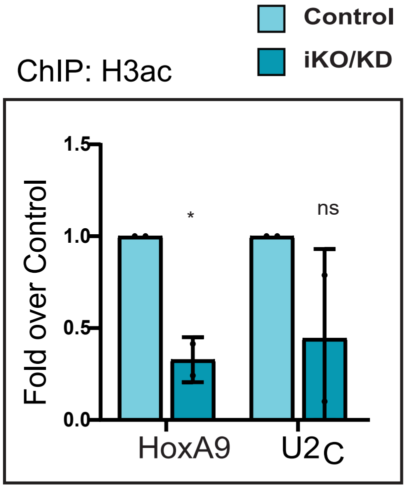


Lone Supplementary Figure 2b

**Supplementary Figure S3. Multiple histone lysine acetyltransferases (KAT) modulate rDNA transcription.**

**A-B.** (**A**) Represents the individual KAT transcript levels upon the shRNA-mediated knockdown of the MOF, MOZ, P300, CBP, PCAF and GCN5, while (**B**) represents the 5'ETS levels upon shRNA knockdown. Data are derived from three biological replicates, with error bars representing the mean ± SD. Statistical significance was assessed using a two-way ANOVA with Sidak's multiple comparison test (*P ≤ 0.05,**P ≤ 0.00005).

**C-J**. The 5'ETS levels upon the overexpression of the PCAF, CBP, P300 and GCN5 are shown (**C-F**),and the transcript levels of the PCAF, CBP, P300 and GCN5 are shown below (G-J). Data are derived from three biological replicates, with error bars representing the mean ± SD.

**K.** Represents the ChIP-seq tracks for the FLAG-MOZ, FLAG-MOZ HAT mutant and H3K9ac ChIP-seq analysis done in the cells overexpressing the vector control; FLAG-MOZ and MOZ HAT mutant are shown. The annotation of an individual human rDNA repeat highlights its various regions, including the intergenic spacer (IGS), spacer promoter (Sp_prom), 47S core promoter, 5' external transcribed sequence (5_ETS), and 3' external transcribed sequence (3_ETS) is shown at the bottom.

**L**. Represents the ChIP-seq tracks for the MOF, H4K16ac, H4K12ac, H4K8ac and H4K5ac in the control and auxin-treated conditions of mouse embryonic stem cells. MOF was acutely depleted using an auxin-inducible degron system here. The annotation of an individual mouse rDNA repeat, as described in S1A, is shown at the bottom.

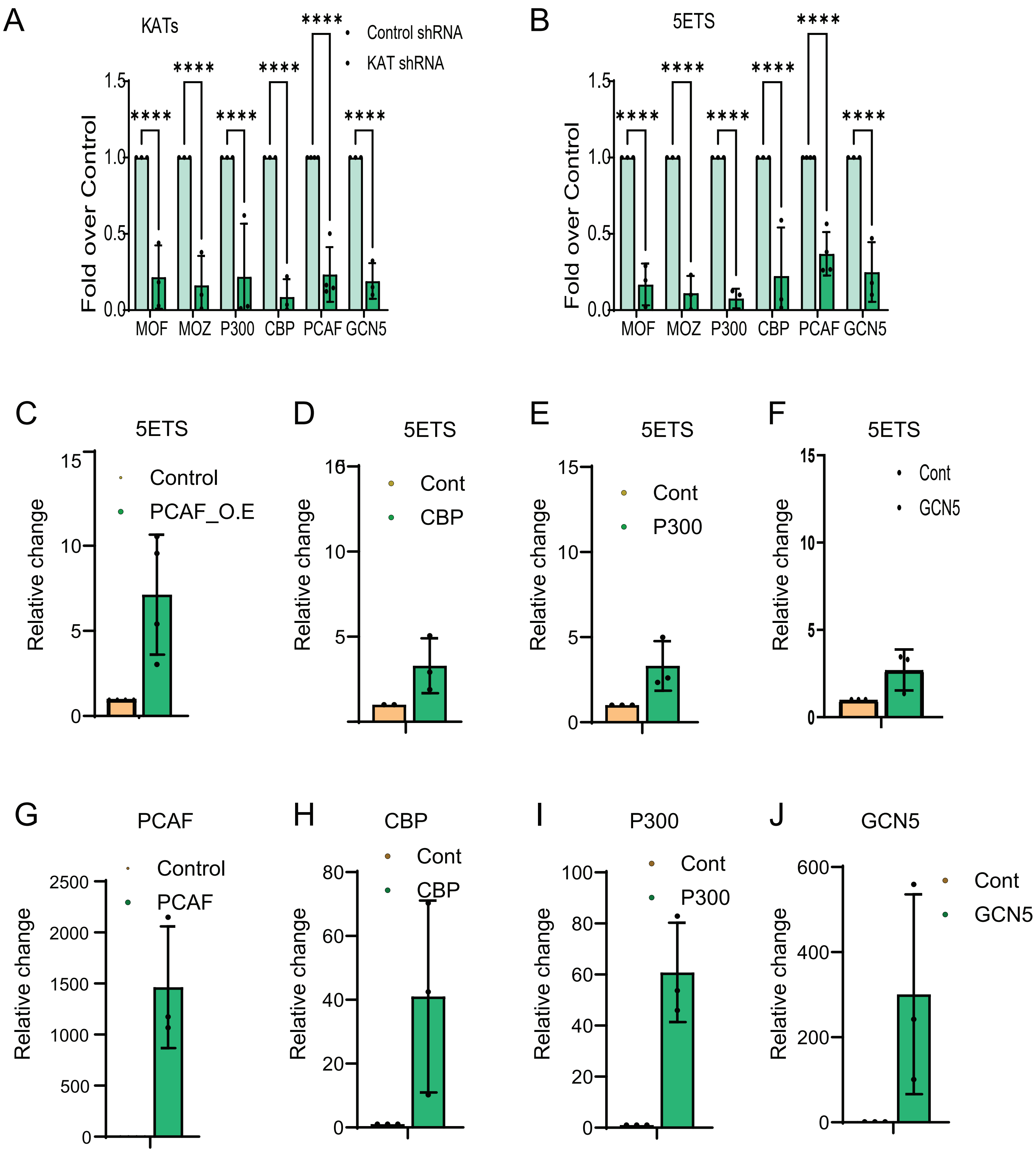


Lone Supplementary Figure 3a

ChIP-seq signals

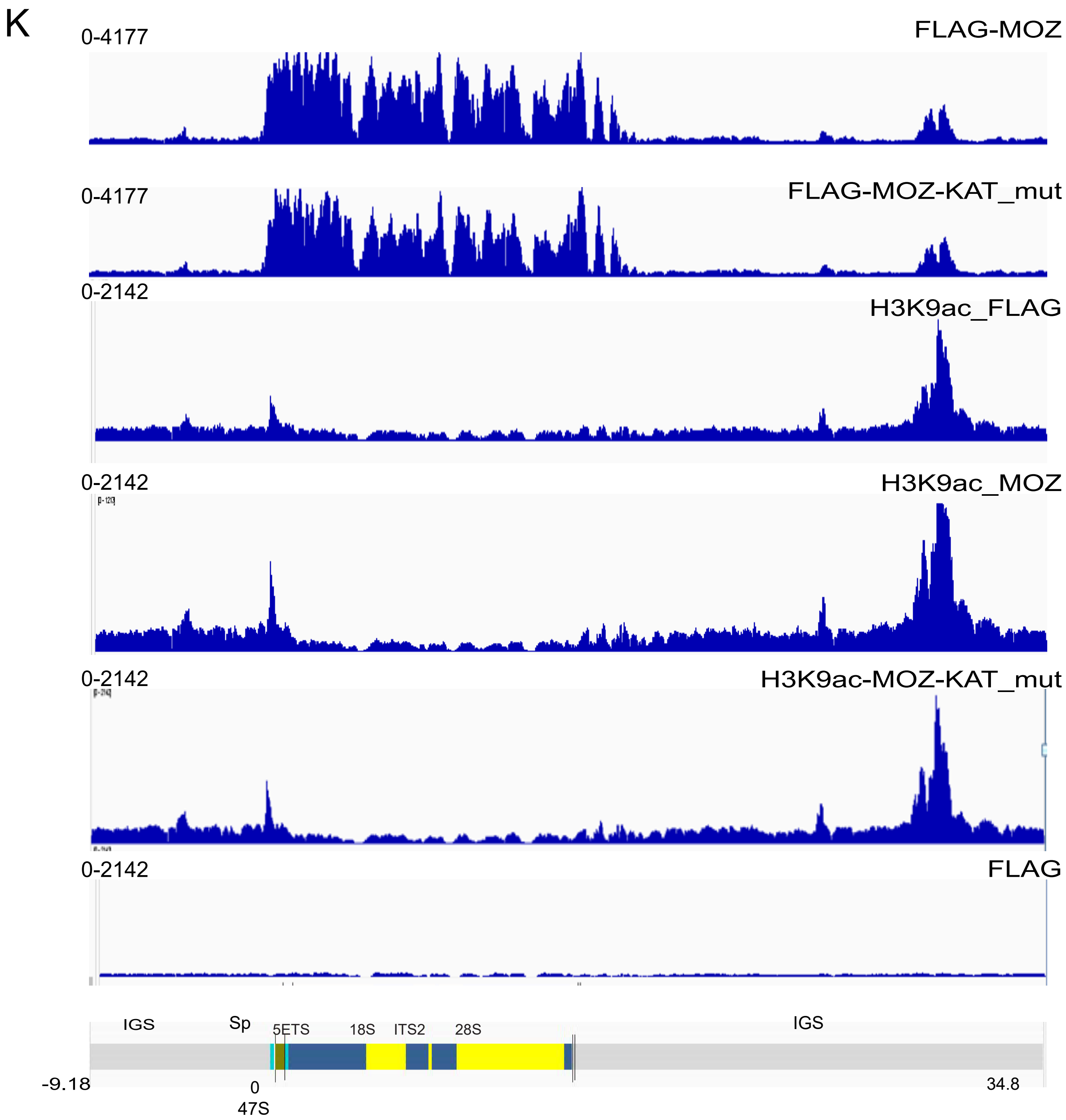

L

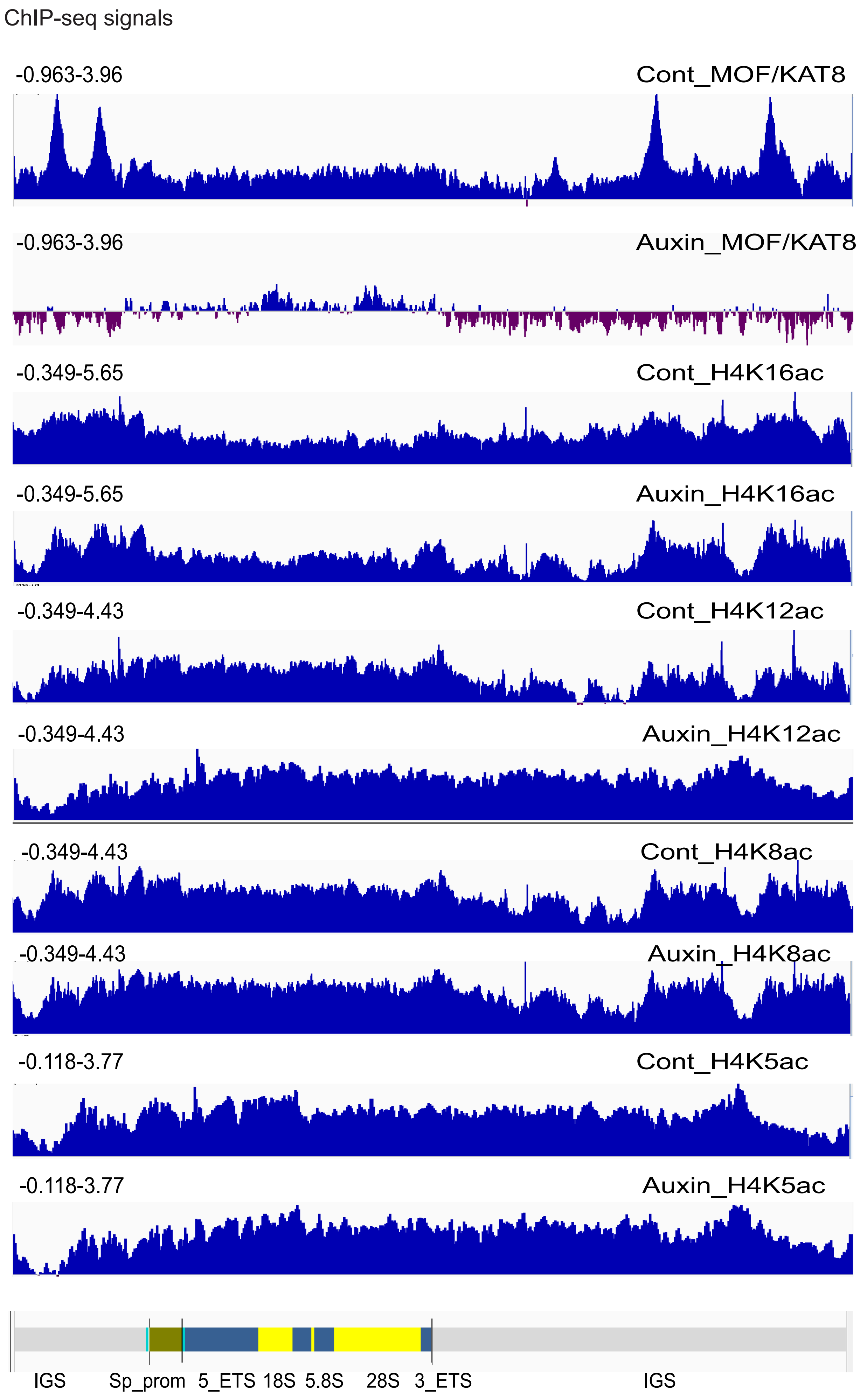


Lone Supplementary Figure 3c

**Supplementary Figure S4.The TA domain of KMT2A regulates rDNA transcripts.**

**A-C**. This figure depicts the transcriptional analysis of KMT2A (**A**), 5′ETS (**B**), and Alpha-satellite (**C**) transcripts in HEK293 cells expressing control, KMT2A, KMT2AΔSET and KMT2AΔTAD. Statistical significance was calculated relative to the control condition in each bar graph. Error bars represent the standard deviation from independent biological replicates. Statistical analysis was performed using a two-way ANOVA with Šídák's multiple comparison test, with $^{*}P \leq 0.05$, $^{**}P \leq 0.005$, and ns (not significant) for $P > 0.05$.

**D**. The immunoprecipitation of endogenous KMT2A in HEK293 cells followed by immunoblot probed with anti- PCAF and KMT2A antibodies is shown. IgG was used as a mock control.

**E**. Ectopically expressed SFB-KMT2A-D1 and SFB alone were subjected to S-protein pull-down assays followed by immunoblot analysis with antibodies against PCAF and FLAG (to detect KMT2A-D1 and SFB) as shown. Molecular weight markers are indicated on the left.

**F**. Represents the ChIP analyses of PCAF in the control and the KMT2A knockout conditions. *HoxA9* and *U2C* were used as the positive and negative control primers. Data are derived from three or more biological replicates, with error bars representing the mean ± SD. Statistical significance was assessed using a two-tailed Student's t-test ($^{*}P \leq 0.05$, $^{**}P \leq 0.005$, ns: not significant, $P > 0.05$).

**G-H**. Shown are KMT2A knockdown levels in cells treated with either Control siRNA or KMT2A-siRNA (G) analyzed for relative KMT2A expression using KMT2A-specific primers. rDNA transcript levels were assessed using different primers: Primer 1 targeted the spacer promoter, Primer 2 targets the enhancer repeats (ER), Primer 3 targeted the 47S promoter, and regions labelled 3' and 3" represent 5'ETS transcript levels in both control and KMT2A knockdown conditions. A schematic representation of these primers is provided below the graph. Both experiments (G-H) were performed in biological triplicate. Error bars indicate standard deviation, and statistical significance was determined using Student's t-test (*$P \leq 0.05$, ns: not significant, $P > 0.05$).

**I**. KMT2AΔSET and KMT2AΔTAD constructs were expressed in KMT2A iKO cells followed by immunoblot analysis with antibodies against KMT2A and α-tubulin as shown. Molecular weight markers are indicated on the left.

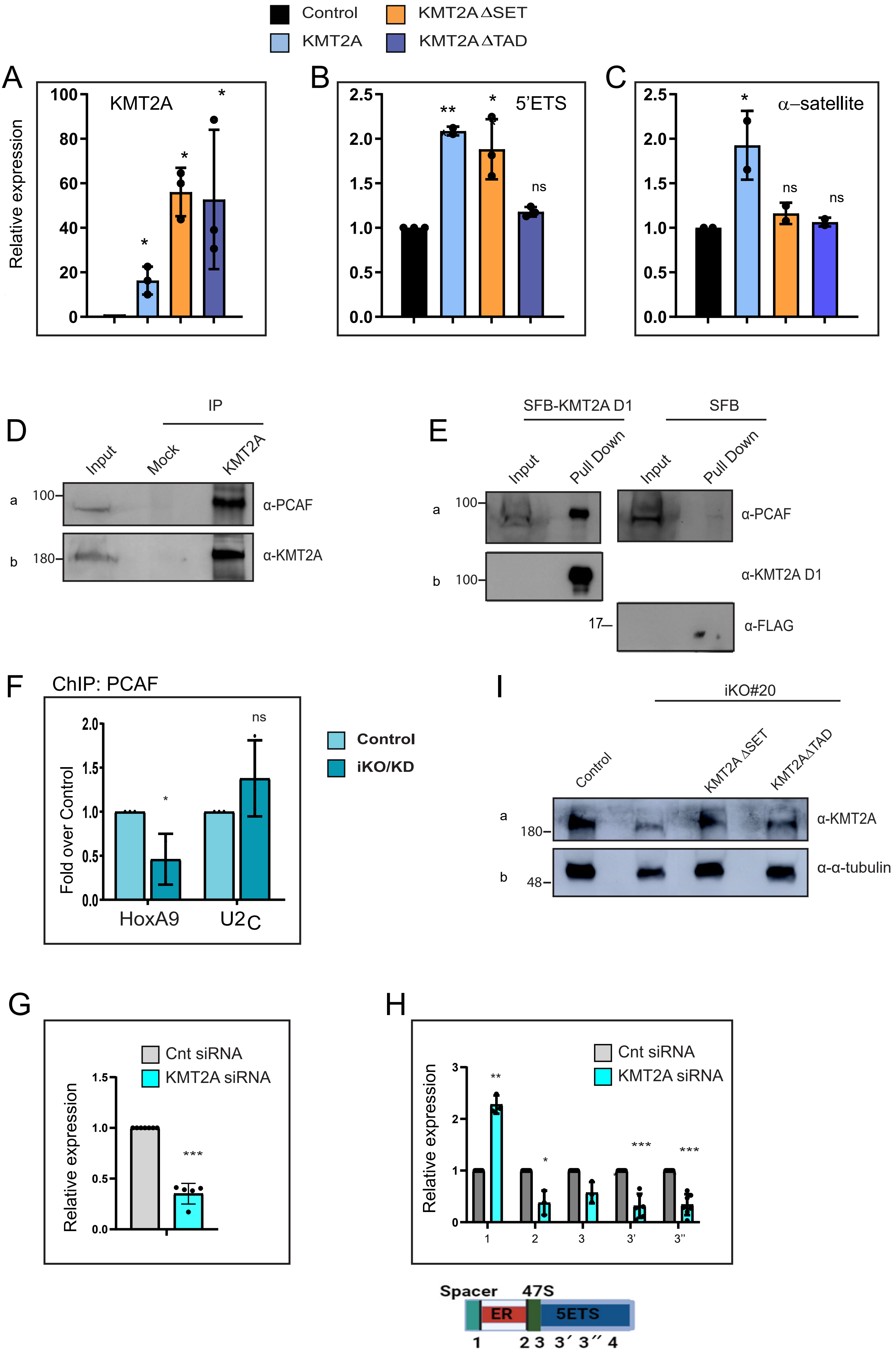


Lone Supplementary Figure 4
