## Supplemental Table for "KMT2A modulates the epigenetic landscape of rDNA by facilitating the recruitment of histone lysine acetyltransferase PCAF to the rDNA locus"

| Primer no. | Position | Primer Sequence (Human rDNA) | Source |
|---|---|---|---|
| 1 | -813 / -707 | F ACATAAACCTGCACGCCAGA | Lone et.al; 2024 |
| | | R CTAGGCAGAGCTCCGGAAA | |
| 2 | -410 / -272 | F GATCCTTTCTGGCGAGTCC | Zenter et al; 2011 |
| | | R GGAGCCGGAAGCATTTTC | |
| 3 | -156 / +43 | F GTGTGTGGCTGCGATGGT | Zenter et al; 2011 |
| | | R CCAACCTCTCCGACGACAG | |
| 4 | +1146 / +1305 | F GGTCGTGTGTGGGTTGACTT | Zenter et al; 2011 |
| | | R GCGGTACGAGGAAACACCT | |
| 5 | +3990 / +4092 | F CGACGACCCATTCGAACGTCT | Zenter et al; 2011 |
| | | R CTCTCCGGAATCGAACCCTGA | |
| 6 | +6697 / +6775 | F GCAGGACACATTGATCATCG | Zenter et al; 2011 |
| | | R GACGCTCAGACAGGCGTAG | |
| 7 | +8256 / +8344 | F GCTAAATACCGGCACGAGAC | Zenter et al; 2011 |
| | | R TTCACGCCCTCTTGAACTCT | |
| 8 | +12855 / +12970 | F ACCTGGCGCTAAACCATTCGT | Zenter et al; 2011 |
| | | R GGACAAACCCTTGTGTCGAGG | |
| 9 | +18449 / +18591 | F TGGTGGGATTGGTCTCTCTC | Zenter et al; 2011 |
| | | R CAGCCTGCGTACTGTGAAAA | |
| 10 | +24,028 / +24116 | F CCCGCGCACATAATAACTAA | Lone et.al;2024 |
| | | R AAATCACTCCTCACGGGAAC | |
| 11 | +28163 / +28298 | F CACTACCCACGTCCCTTCAC | Zenter et al; 2011 |

| | | | |
|---|---|---|---|
| | | R GAGAGAAGACGGAGGCACAC | |
| 12 | +28279 / +28456 | F GTGTGCCTCCGTCTTCTCTC | Zenter et al; 2011 |
| | | R GTCAAGGGGCTATGCCATC | |
| 13 | +28328 / +28495 | F ATTCTTGCCAGGCTGACATT | Zenter et al; 2011 |
| | | R AAGCCTCACAACTGCAGACC | |
| 14 | 28475 / +28597 | F GTCTGCAGTTGTGAGGCTTT | Lone et al; 2024 |
| | | R CGGAGGGCGGAGAAACTAAA | |
| 15 | +30541 / +30640 | F ACTGGCGAGTTGATTTCTGG | Zenter et al; 2011 |
| | | R CGAGACAGTCGAGGGAGAAG | |
| 16 | +37,995 / +38,09 | F CTCACAGAGGAAGGGAGCAC | Lone et al; 2024 |
| | | R AACAAGGGAGGGAGGAACTT | |
| 17 | +41907 / +42035 Or - 1092 / - 964 | F CCGTGGGTTGTCTTCTGACT | Zenter et al; 2011 |
| | | R AAGCGAAACCGTGAGTCG | |
| 18 | +42012 / +42202 | F GCTTCTCGACTCACGGTTTC | Zenter et al; 2011 |
| | | R GGAGCTCTGCCTAGCTCACA | |
| **Canonical Gene Primers** | | | |
| *HoxA9* | Fw. | CTCCGCCGCTCTCATTCTCAG | Malik et.al; 2022 |
| | Rev. | GCCAGAAGGGGTGACTGTCC | |
| *Rad18* | Fw. | ATGCGCAGTACAAGCCCTTA | Malik et.al; 2022 |
| | Rev. | GCTCCAACACCACTCGAAAT | |
| U2C | Fw. | TTTGCTCCCACTGCCGTC | Malik et.al; 2022 |
| | Rev. | CTGAGTCTTTCGGTGCCC | |

| | | | |
|---|---|---|---|
| CD4 | Fw. | TCTGCAGAAGGAACAAAGCA | Malik et.al; 2022 |
| | Rev. | GGAAGGAAGCCGAGTCTGA | |
| **Transcriptional Primers** | | | |
| *MLL1* | Fw. | GGAGCACACATTCCAGACCA | Malik et.al; 2022 |
| | Rev. | TTTGGGTCACCTGAACTTCC | |
| *SETD1A* | Fw. | CGAATACGTGGGTCAGAACA | Malik et.al; 2022 |
| | Rev. | TGCAGCAGTGGTTGATGAAT | |
| 3′ | Fw. | GTCCCCTCGTCTGCTCCTCTC | Lone et al; 2024 |
| | Rev. | CAAGTCGACAACCACTG | |
| 3″ | Fw. | CCTGCTGTTCTCTCGCGCGTCCGAG | Lone et al; 2024 |
| | Rev. | AACGCCTGACACGCACGGCACGGAG | |
| **Mouse rDNA Primers** | | | |
| 1. | Fw. | CGACCAGGGTGACAGGAG | This study |
| | Rev. | GCTTTGGCCATCTCCCTGTA | |
| 2. | Fw. | GAAGCCCTCTCTGTCCCTG | This study |
| | Rev. | ACCCGGAGAACTGATAAGACC | |
| 3. | Fw. | GGTTCTTTTCGTTATGGGGTCA | This study |
| | Rev. | AGCATAAAAGAGACAGGGAGGA | |
| 4. | Fw. | CTTGCGTGTGCTTGCTGT | Zenter et.al; 2013 |
| | Rev. | GAAATCGGGAAAAACGTCTG | |
| 5. | Fw. | TGTCTGCCCGTATCAGTAACTGTC | Zenter et.al; 2013 |
| | Rev. | CCCTGGCCCGAAGAGAACT | |
| 6. | Fw. | CATCTGCTCTGGTCGAGGTT | Zenter et.al; 2013 |

| | | | |
|---|---|---|---|
| | Rev. | GCAAGACCCAAACACACACA | |
| 7. | Fw. | GGAATCCGATGCACACTTTT | Zenter et.al; 2013 |
| | Rev. | TGTGTGTGTGTGTGTGTGTGA | |
| 8. | Fw. | AACTGTGCCTGTTCCCTCAC | Zenter et.al; 2013 |
| | Rev. | GGCACCCAAAAACGAAAGTA | |
| 9. | Fw. | GACACAGGAGAGGGAAGTGC | Zenter et.al; 2013 |
| | Rev. | CTCCCTGTACGACCTCCTTG | |
